# Hyperdiverse, bioactive, and interaction-specific metabolites produced only in co-culture suggest diverse competitors may fuel secondary metabolism of xylarialean fungi

**DOI:** 10.1101/2024.12.10.624342

**Authors:** Mario E.E. Franco, Megan N. Nickerson, Benjamin P. Bowen, Katherine Louie, Trent R. Northen, Jana M. U’Ren

**Author notes:** Sustainable Plant Protection Programme, Institute of Agrifood Research and Technology (IRTA), Lleida, Spain.

## Abstract

Xylariales is one of the largest and most ecologically diverse fungal orders that is well-known for its chemical diversity. Enhanced secondary metabolism of Xylariales taxa is associated with increased gene duplication and horizontal gene transfer (HGT) of biosynthetic gene clusters (BGCs), especially in generalist taxa with both greater saprotrophic abilities and broader host ranges as foliar endophytic symbionts. Thus, one hypothesis for BGC diversification among more generalist fungi is that diverse competitive interactions—in both their free-living and symbiotic life stages with many hosts—may exert selective pressure for HGT and a diverse metabolic repertoire. Here, we used untargeted metabolomics to examine how competition (pairwise co-cultures) between seven xylarialean fungi influenced their metabolite production. Of the >9,000 total features detected, 6,115 and 2,071 were over-represented in co-cultures vs. monocultures, respectively. For each strain, each additional co-culture interaction resulted in an 11-14-fold increase in metabolite richness compared to monocultures, reflecting the limited amount of metabolite overlap among different co-culture combinations. Phylogenetic relatedness and BGC content did not impact the diversity of metabolites produced in co-culture; however, co-cultures between more ecologically distinct fungi elicited the strongest metabolic response. Overall, the diversity, specificity, and putative bioactivity of metabolites over-represented only in co-culture supports the role of widespread and diverse competitive fungal interactions to drive xylarialean metabolic diversification. Additionally, as fungal-produced plant hormones were only detected in co-culture, our results reveal the potential for *in planta* interactions among fungal endophytes to influence the host plant.

**Importance:** Saprotrophic and endophytic xylarialean fungi are among the most prolific producers of bioactive secondary metabolites, with numerous industrial uses as antibiotics, pharmaceuticals, and insecticidal toxins. Fungal secondary metabolites are typically encoded in biosynthetic gene clusters (sets of physically clustered genes), but the products of most clusters are unknown as the genes are not active in typical culture conditions. Co-cultures can help to “turn on” fungal secondary metabolite production, yet factors that can influence co-culture outcomes are largely unknown. Here, we used untargeted metabolomics to assess how differences in genomic content, ecology, and phylogenetic relatedness among seven diverse xylarialean fungal strains impact metabolic production in co-culture. As expected, co-culturing significantly increased metabolite diversity, as well as the abundance of putatively bioactive metabolites. Each new pairwise combination produced different metabolites, indicative of strain-specific responses to competitors. This new information will enable further characterization of the immense biotechnological potential of xylarialean fungi.

## Introduction

As both plant symbionts and saprotrophs that can catabolize a wide range of organic substrates, fungi are a critical component of all ecosystems (1, 2). As osmotrophs, competition for shared resources can drive fungal metabolic diversification, especially the production of secondary metabolic (SM) compounds that provide an ecological advantage (3, 4). Fungal SMs can determine virulence against plants and animals, enhance nutrient acquisition and defense against predators and abiotic stressors, and facilitate intra-organismal communication and quorum sensing (e.g., (4–7)). Genes involved in fungal SM production are frequently encoded by biosynthetic gene clusters (BGCs) that can evolve through gene duplication and/or horizontal gene transfer (HGT) (e.g., (8, 9)).

Xylarialean fungi, especially species of Xylariaceae *sensu lato* (*s.l*.) (10, 11), are among the most prolific fungal SM producers (12, 13). Described species are primarily saprotrophs or pathogens of woody hosts (14–16); however, numerous species have been cultured as endophytic symbionts of diverse plants and lichens (e.g., (17–19)). Endophytic xylarialean fungi appear to be host and substrate generalists: closely related strains have been cultured from phylogenetically diverse hosts, as well as both living and decomposing plant tissues (17). A previous study found that xylarialean endophytes with higher BGC diversity and rates of HGT formed symbiotic associations with a greater diversity of hosts and displayed a greater capacity to degrade leaf litter (11). Generalist endophytes thus interact with both a greater number of hosts, as well as potentially with diverse microbes *in planta* and/or during a free-living saprotrophic stage, which may promote greater SM diversification compared to more ecologically specialized species (11) (see also (20)).

Here, to better understand the evolutionary and ecological factors influencing SM diversification in xylarialean fungi, we used untargeted metabolomics to study the metabolic interactions in all pairwise combinations of seven xylarialean fungal strains whose BGC diversity was previously studied (11) (Fig. 1A). Although performed *in vitro*, co-culturing can induce the production of bioactive compounds not typically produced by either microorganism when grown without competitors (21–24). For xylarialean fungi, co-culture has proven to be a valuable tool to trigger the production of bioactive SMs (25–28). More recently, co-culturing paired with additional techniques such as mass spectrometry, NMR, and structural elucidation has led to discoveries of novel SMs with applications in the pharmaceutical, food, and agronomic industries (21, 23, 29).

**Figure 1.**
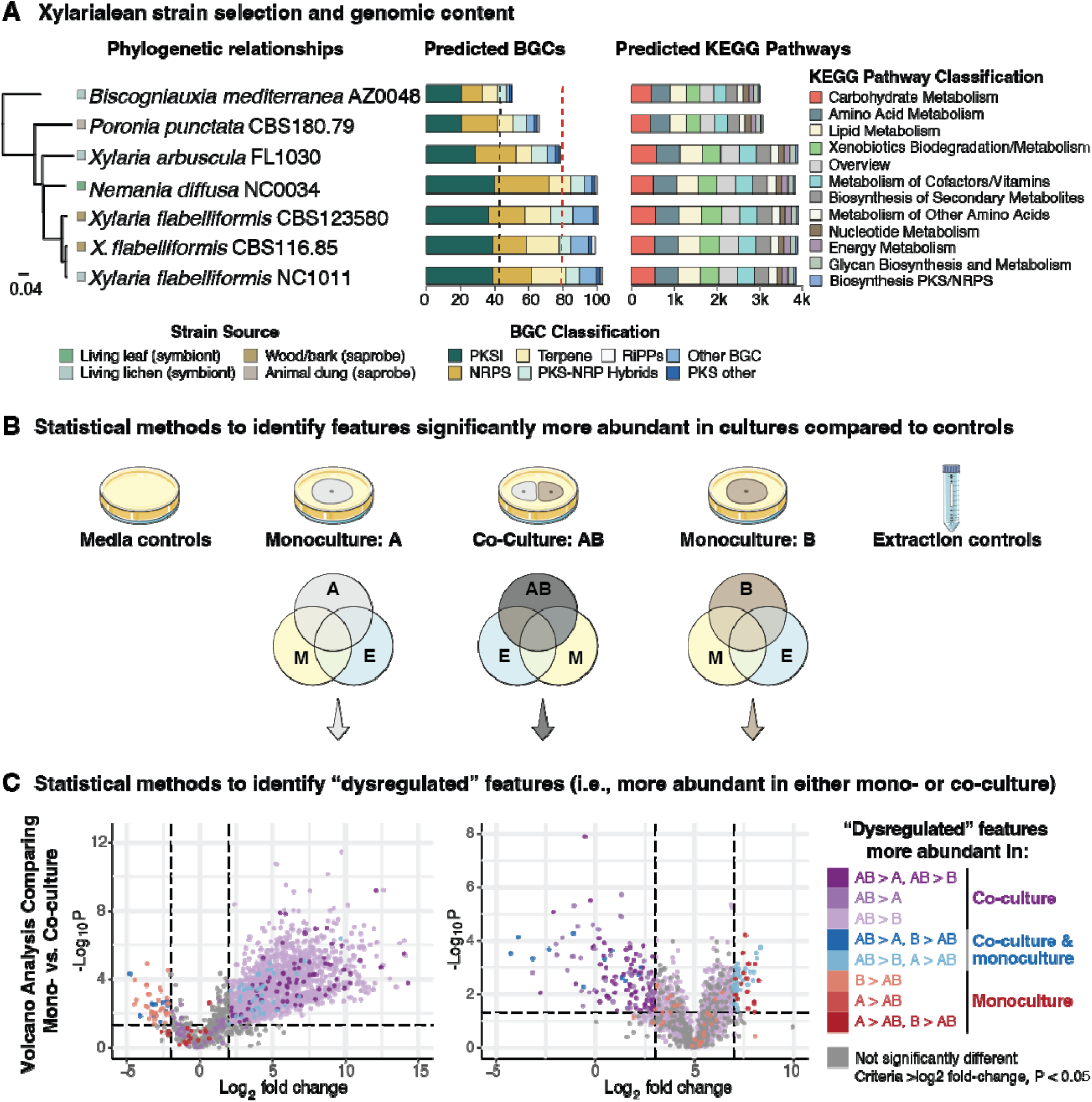
Ecology, phylogeny, and genomic content of seven xylarialean strains selected for study and the statistical analysis workflow used to identify metabolite features significantly more abundant in co-culture or monoculture for 21 different pairwise comparisons. (**A**) Phylogenetic relationships and ecological mode of the seven strains, which included three closely related strains of *Xylaria flabelliformis,* along with four additional strains of Xylariaceae *s.l.* that were increasingly phylogenetically distant from *X. flabelliformis.* Different colored boxes next to each strain indicate its substrate of isolation and trophic mode (symbiont or saprobe). Stacked bars indicate the number and classification of predicted BGCs (lines indicate the mean number of BGCs in genomes of other Pezizomycotina (black) vs. Xylariaceae *s.l.,*(red)) and abundance of different KEGG pathways in each strain (see Supplemental Tables 7-9) (11). (**B**) Schematic of statistical methods to identify features from each monoculture (above A or B) and co-culture (AB) whose abundance was log2FC greater than controls (i.e., three media-only plates (extracted in parallel) and extraction controls). In total, there were three replicate plates for each pairwise co-culture (n = 63), each of the seven monocultures (n = 21) for a total of 84 samples. (**C**) Example of the pairwise univariate volcano plot analyses used to identify features significantly over-abundant in either co-culture or monoculture (“dysregulated”; criteria >log2 fold-change (log2FC), p-value < 0.05). As illustrated, features from each co-culture (AB) were compared to each corresponding monoculture (A or B) and assigned significance in either treatment: monoculture (red colors) or co-culture (purple colors) (see Table S1). Given the nature of the pairwise comparisons, some features (blue colors) could not be unambiguously assigned to either category: they were more abundant in co-culture compared to at least one of the monocultures, but also more abundant in at least one of the monocultures compared to a co-culture (i.e., AB > B and A > AB; AB > A and B > AB; hereafter, “ambiguous” features) (see Table S2). Data from both chromatography columns and ionization modes were combined for analyses (with duplicates removed).

Our main goals were to statistically compare the diversity, composition, and bioactivity of small molecules secreted by fungi in monoculture vs. co-culture, explicitly quantifying the impact of fungal ecological mode, BGC content, and phylogenetic position on the resulting metabolome diversity and composition. Specifically, we tested three hypotheses: (i) a strain’s BGC content will be positively correlated with the number of metabolites it produces; (ii) the phylogenetic distance between competing strains will influence metabolite production, with greater competition and SM production occurring in interactions between close relatives (21, 22); and (iii) co-culture interactions will yield a greater number and diversity of metabolites compared to monocultures, with fungi exhibiting strain-specific metabolic responses to different competitors rather than producing a uniform suite of metabolites for all competitors. As our primary goal was to detect bioactive SMs involved in competition, here we focused on the metabolites secreted into the interaction zone between cultures rather than the entire extra- and intercellular metabolome.

## Results

### Fungal interactions

Overall, 17 of 21 (80%) pairwise co-culture combinations resulted in deadlock growth between strains (i.e., fungal hyphae approached each other, but growth was arrested). Deadlock usually resulted in a clear space between fungal hyphae representing a potential zone of inhibition, but in some co-cultures, hyphae of each pair appeared to nearly touch, yet without a clear zone (Supplemental Figs. 1-7). Only four co-culture pairs resulted in the invasion of one strain over another: three cases involved *X. arbuscula* invading other strains, and the fourth instance was *B. mediterranea* overgrowing *P. punctata*.

### Results of Untargeted Metabolomics

Across all 84 fungal samples and controls, liquid chromatography coupled with high-resolution tandem mass spectrometry (LC-MS/MS) detected 47,316 ions with unique mass-to-charge ratio (m/z) and retention time (RT) values (i.e., features) (Table S1). Approximately 20% of features (n = 9,762) were significantly more abundant in fungal cultures compared to negative extraction and media controls (Table S1; Table S3). Only 955 of 9,762 features were assigned to a Superclass (Fig. S8).

### Influence of co-culture vs. monoculture on metabolite production

Pairwise statistical analyses of all 21 co-cultures compared to their respective monocultures (Fig. 1B) identified a total of 6,141 features that were significantly over-represented in either co-culture and/or monoculture (hereafter, “dysregulated”) (Table S2). Among these dysregulated features, 4,070 were unique to co-cultures (i.e., only over-represented in co-culture), and 26 features were unique to monocultures (i.e., only over-represented in monoculture). Including features over-abundant in at least one co-culture or monoculture (i.e., “ambiguous”; see Fig. 1B), our pairwise analyses identified a total of 6,115 features that were significantly more abundant in at least one co-culture vs. monoculture. In comparison, 2,071 features were more abundant in at least one monoculture vs. its respective co-cultures (Table S2).

The mean number of features significantly more abundant in at least one co-culture was >2-fold higher than the mean number of features significantly over-represented in at least one monoculture (Fig. 2A; Table S2). Moreover, rarefaction analysis illustrates each additional different co-culture interaction increased metabolite richness (i.e., the number of metabolites) 11 to 14-fold (i.e., 1,279 to 4,037 additional metabolites) compared to similar comparisons between monocultures (Fig. 2B). Although the number of features over-represented in co-cultures did not differ significantly between fungi of various phylogenetic distances (Fig. 2C), co-cultures involving *P. punctata* resulted in a 1.8X increase in the mean number of over-represented features regardless of partner (Fig. 2D).

**Figure 2.**
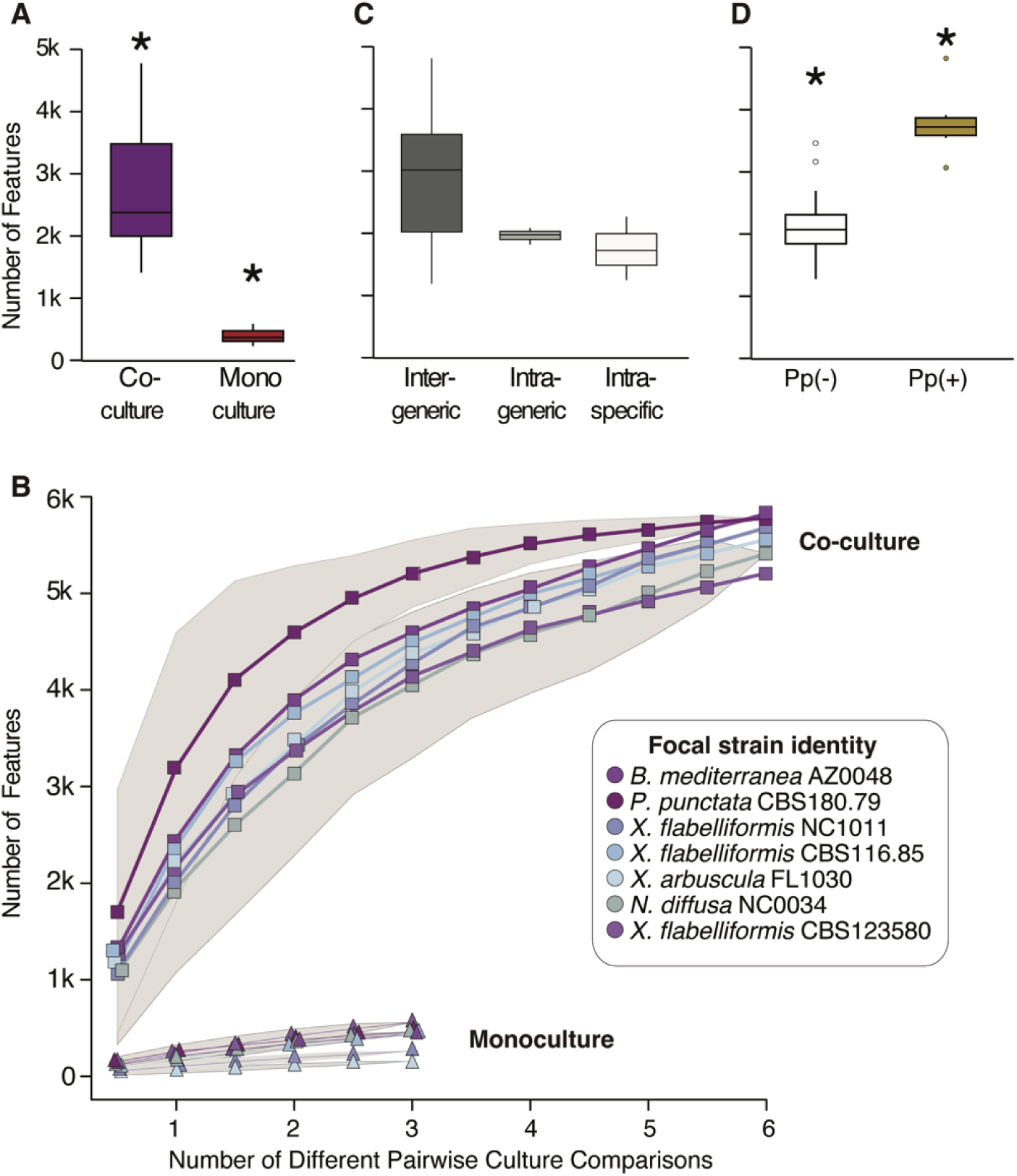
Each additional co-culture combination results in a significant increase in metabolite richness compared to monocultures, but the magnitude of metabolite production does not reflect phylogenetic relationships. (**A**) Interquartile box plots comparing the median number of significantly over-abundant metabolites produced in all pairwise co-culture comparisons vs. all monoculture comparisons. Asterisks indicate significant differences after t-test of means (t_20.6_ = 11.56, P <0.0001). (**B**) Rarefaction curves of metabolite richness as a function of the number of comparisons to other cultures when grown in either co-culture or monoculture. (**C-D**) Interquartile box plots comparing the median number of metabolites that were significantly over-abundant in co-culture as a function of the taxonomic relationships among different pairs. (**C**) Intergeneric: pairwise comparisons of strains in four different genera (*Xylaria*, *Nemania*, *Poronia*, and *Biscognauxia*); Intrageneric: pairwise comparisons of four *Xylaria* spp.; Intraspecific: pairwise comparisons of three *X. flabelliformis* strains). ANOVA found no significant differences: F_2,18_ = 2.60, P = 0.1000. (**D**) Pairwise co-culture comparisons either with (Pp+) or without (Pp-) strain *P. punctata* CBS 180.79. Asterisks indicate significant differences after t-test of means (t_9.5_ = −5.84, P = 0.0002).

Although co-cultures consistently resulted in a greater number of over-represented features compared to monocultures, the magnitude of the difference was variable among different pairwise comparisons (Fig. 3). For example, we observed a 6-fold increase in significant features for the co-culture of *P. punctata* and *X. flabelliformis* NC1011 vs. either strain’s monoculture. However, there was only a 1.2X increase in the number of features for the co-culture of *X. arbuscula* vs. *N. di usa* compared to either monoculture (Table S2). The latter pair also had the fewest total features in co-culture (n = 1,278). The highest number of features significantly over-represented in any co-culture was 4,817 (*P. punctata* vs. *X. flabelliformis* CBS 116.85), whereas the highest number of features over-represented in monoculture compared to co-culture was 405 for *N. di usa* NC0034 vs. *B. mediterranea* AZ0048 (Table S2). (Table S2).

**Figure 3.**
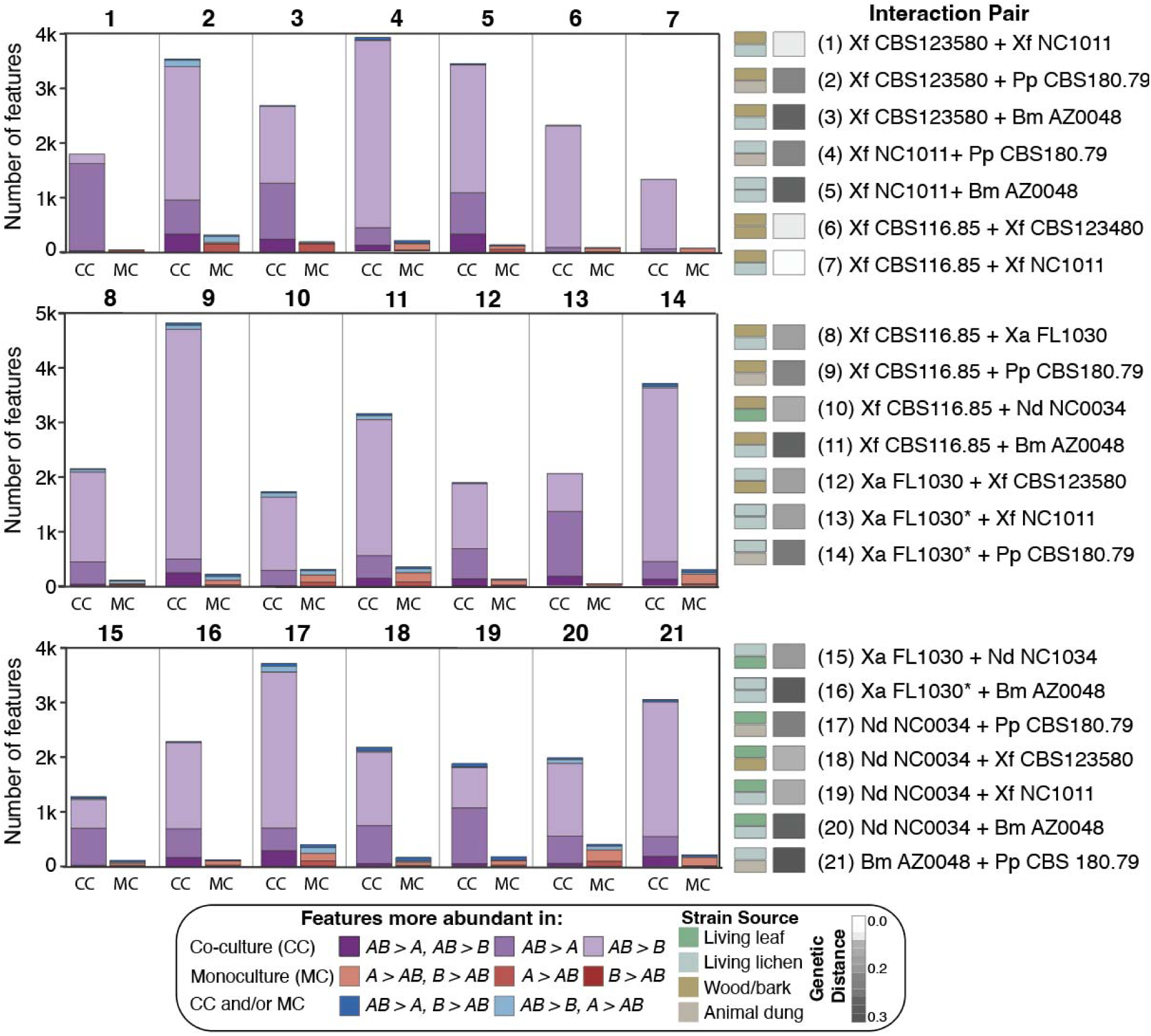
Co-culture results in the production of significantly more features relative to monocultures, yet the magnitude of the increase varies across 21 different pairwise strain combinations. Stacked bar plots with the total number of features that were identified as significantly more abundant in either co-culture (CC) or monoculture (MC) for all 21 interaction pairs. Bars are colored by the statistical methods used to assign features to either treatment, as shown in Fig. 1C. Ambiguously defined features (defined as CC and/or MC) are also indicated. Each numbered plot corresponds to one co-culture interaction pair (abbreviated strain names are listed to the right). Asterisks (*) next to *X. arbuscula* (Xa FL1030) indicate three cases where this strain overgrew its competitor in co-culture. For each interaction pair, boxes to the left of the name indicate either the substrate source of each strain or the pairwise genetic distance between genomes of each strain (11). Metabolites were compared after one month of growth either in co-culture or monoculture on *Aspergillus* minimal medium at room temperature under ambient light/dark conditions. Features derived from both chromatography columns and ionization modes were combined into a single dataset and analyzed together (Table S3).

### Influence of competitor identity on co-culture metabolism

Among the 6,115 features that were over-abundant in co-culture vs. the respective monocultures, none were universally over-abundant in all 21 co-culture combinations, and only one feature was shared among 20 of 21 pairs (Fig. 4A). Instead, we found that only 8.4% of over-abundant features (n = 514) occurred in at least 15 of 21 interactions, and only 30.9% (n = 1,891) of features were shared across 11-14 interaction pairs. Instead, the majority of features (60.7%; n = 3,710) were shared among 10 or fewer interactions. In addition, 245 (4.0%) over-abundant features were unique to a single co-culture pair, including 190 features that were unique to the co-culture of *X. flabelliformis* NC1011 and *B. mediterranea* (Fig. 3A). Concomitant with the general lack of shared metabolites among different fungal interactions, a heatmap of feature frequency in co-cultures of each strain illustrates that the production of most metabolite features occurred infrequently (Fig. 4B). Yet, while most features were rare, a subset of features were consistently observed in the co-cultures of some strains (e.g., 533 features that occurred in all co-cultures with *P. punctata* as a partner) (Fig. 4B).

**Figure 4.**
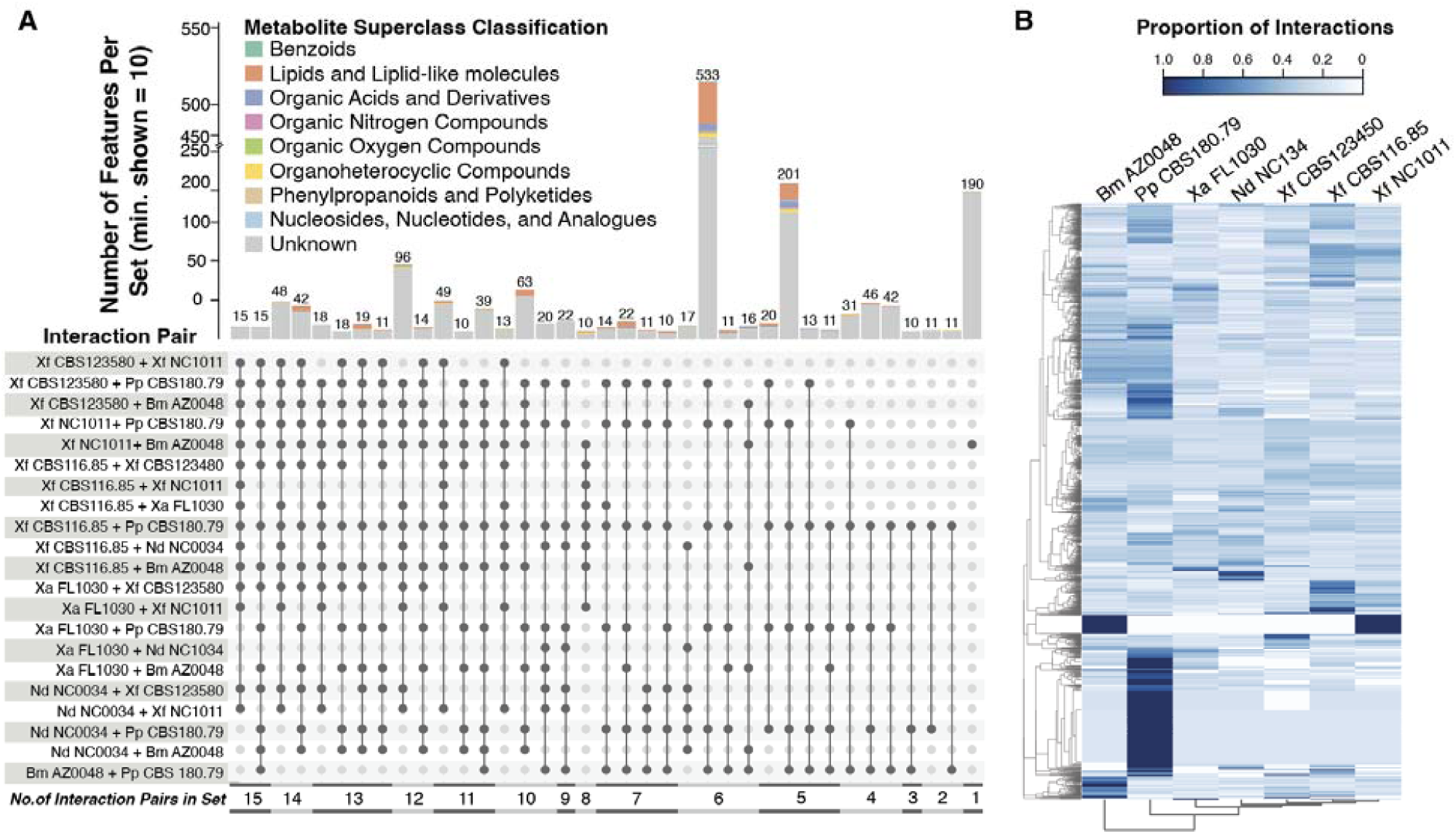
Lack of metabolite overlap across the 21 co-culture pairs is consistent with competitor-specific metabolic responses. **(A)** UpSet plot showing the distribution of features that were over-abundant in co-culture vs. monocultures across all 21 interaction pairs. The height of each stacked bar indicates the number of features found (i.e., “intersection” size) for different interaction pair(s). Vertical black lines connect features shared between >1 pair. Dot color indicates whether a feature is present (black) or absent (grey) for each interaction pair. Colors in each stacked bar correspond to feature classifications within each set at the ‘Superclass level’ using the ClassyFire. Interactions with <10 features are not displayed for readability. **(B)** Heatmap based on the proportion of co-cultures (columns) where a feature (row) was found in co-culture for a particular strain (i.e., number of occurrences out of six different co-cultures). A proportion equal to 1.0 (dark blue) thus corresponds to compounds always detected in co-cultures with that strain, suggesting it may be the compound producer. Compounds and fungal strains were grouped using hierarchical clustering (complete) using Euclidean distance (see also Fig. S11, where strains are instead ordered by their phylogenetic relationships). Features derived from both chromatography columns and ionization modes were combined into a single dataset and analyzed together with duplicates removed.

Similar to the general lack of overlap among features over-abundant in co-culture compared to monoculture across all 21 interactions, comparisons of overlap among features over-represented in six co-cultures of a focal strain compared to its monoculture revealed a majority of features occurred in only one or two co-culture combinations per focal strain, although the percentage of shared features varied by focal strain (Fig. S9). The fraction of features unique to one particular interaction pair ranged from 620 in all co-cultures with *P. punctata* (12% of total features in this strain) to 1,428 in all co-cultures with *X. flabelliformis* NC1011 (53% of total features in this strain). In general, co-cultures with any of the three *X. flabelliformis* strains resulted in a relatively high number of features specific to these interaction pairs (Fig. S9). However, the same 880 features were found among *X. arbuscula* co-cultures with either two (n = 432) or three (n = 448) of the *X. flabelliformis* strains (Fig. S9). Interactions of *B. mediterranea* with all strains except *P. punctata* resulted in the production of the same 686 features (18% percent of 3,776) (Fig. S9). The largest metabolite overlap was observed for interactions with *P. punctata*: 1,096 features were shared among all *P. punctata* co-culture combinations, regardless of the partner (Fig. S9).

### Links between metabolites and fungal genomes

Using MAGI 1.0 (30) we identified 135 features (out of 6,141 total) that were linked to genes/BGCs in the genomes of all seven strains (Table 1). Overall, these 135 features were linked to 2,651 genes via 487 different InChIKey strings (Table 1; Supplemental Tables 4-5). Additionally, 14.3% of linked genes (i.e., 379 genes that were linked to 83 features) were previously found to reside within the boundaries of 300 different BGCs (based on data from (11)). Among these 379 genes, 78 were within BGCs with at least one significant hit against the MIBiG repository (Table 1) for toxic metabolites such as alternariol, asperlactone, cyclosporin, aspirochlorine, etc. (see Table S5) (11).

**Table 1.**
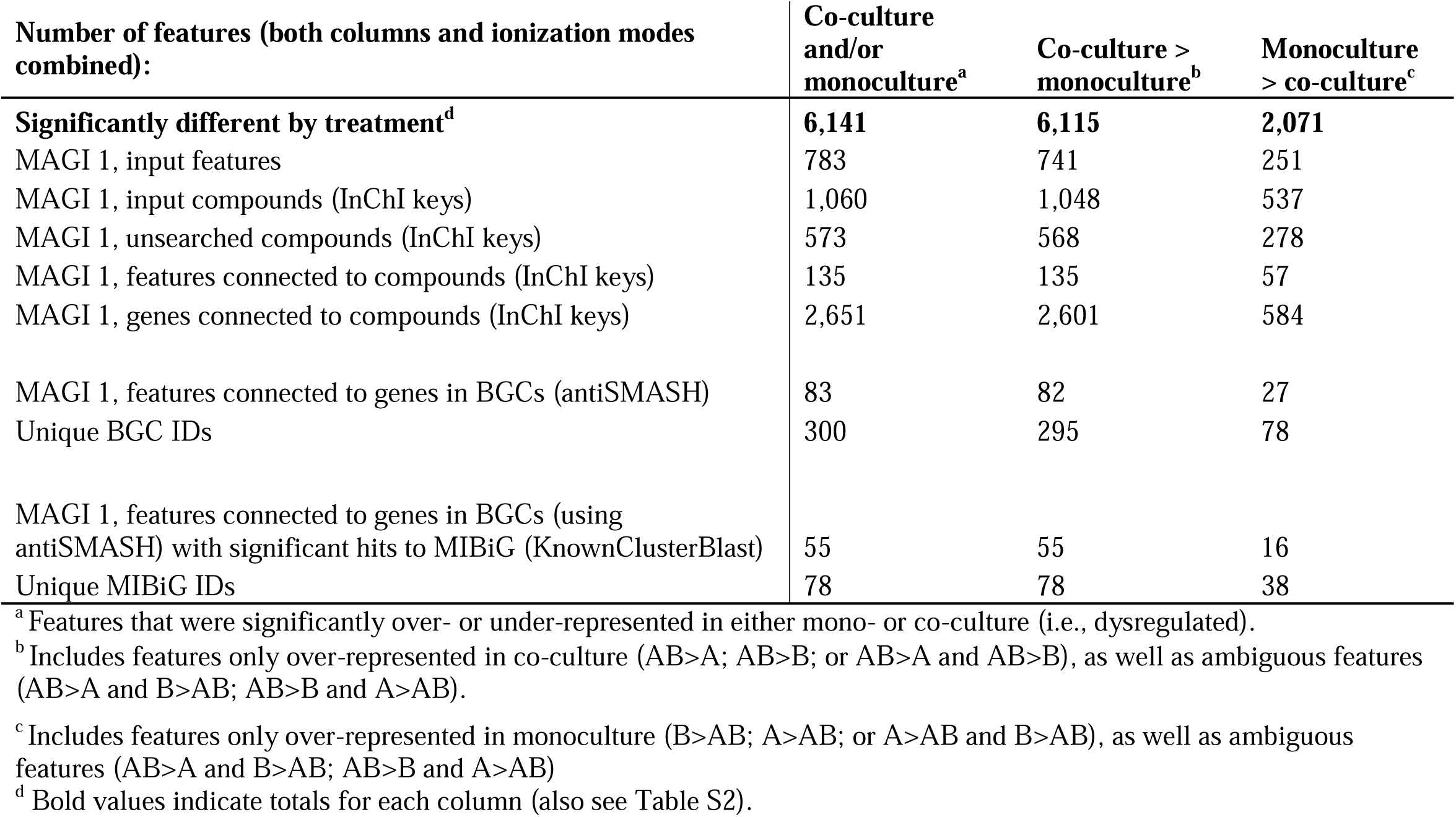
Number of features and related compound-gene connections detected using MAGI 1.

Furthermore, by leveraging previous analyses of HGT in the genomes of these strains (11), we found that 20 of genes MAGI linked to metabolites were predicted to have been acquired via HGT from Bacterial donors (Table S5). Notably, in addition to annotations within BGCs, these 20 HGT candidate genes were predicted to encode enzymes with diverse activities (e.g., GO Terms for catalytic activity, oxidoreductase activity, and transferases). For example, 8 of these 20 genes were annotated as putative opine dehydrogenases (IPR003421:1), which are known to be involved in crown gall tumors produced by plant pathogenic *Agrobacterium* spp. (31).

### Co-culturing and the production of bioactive compounds

In addition to the greater number of features produced in co-cultures vs. monocultures, numerous features that were over-abundant in co-cultures were putatively identified as bioactive compounds (Table 2; Table S6). Many of these features were also linked to genes within BGCs with MIBiG hits to numerous bioactive compounds (see above; Table S5). In general, sesquiterpenes were one of the most prevalent bioactive compound classes produced by these seven fungal strains in co-culture (Table 2). Feature hits include matches to parthenolide (32), alantolactone (33), polygodial (34), zerumbone (35), steviol (36), valerenic acid (37), and nootkatone, which have been reported to have anti-microbial, anti-fungal, insecticidal, anti-inflammatory activity. Numerous features also had hits to mycotoxins such as alternariol, diacetoxyscirpenol, culmorin, and the cytochalasins C, D, and J (38, 39), that all pose a serious threat to human health and food safety (12, 13, 21–23).

**Table 2.**
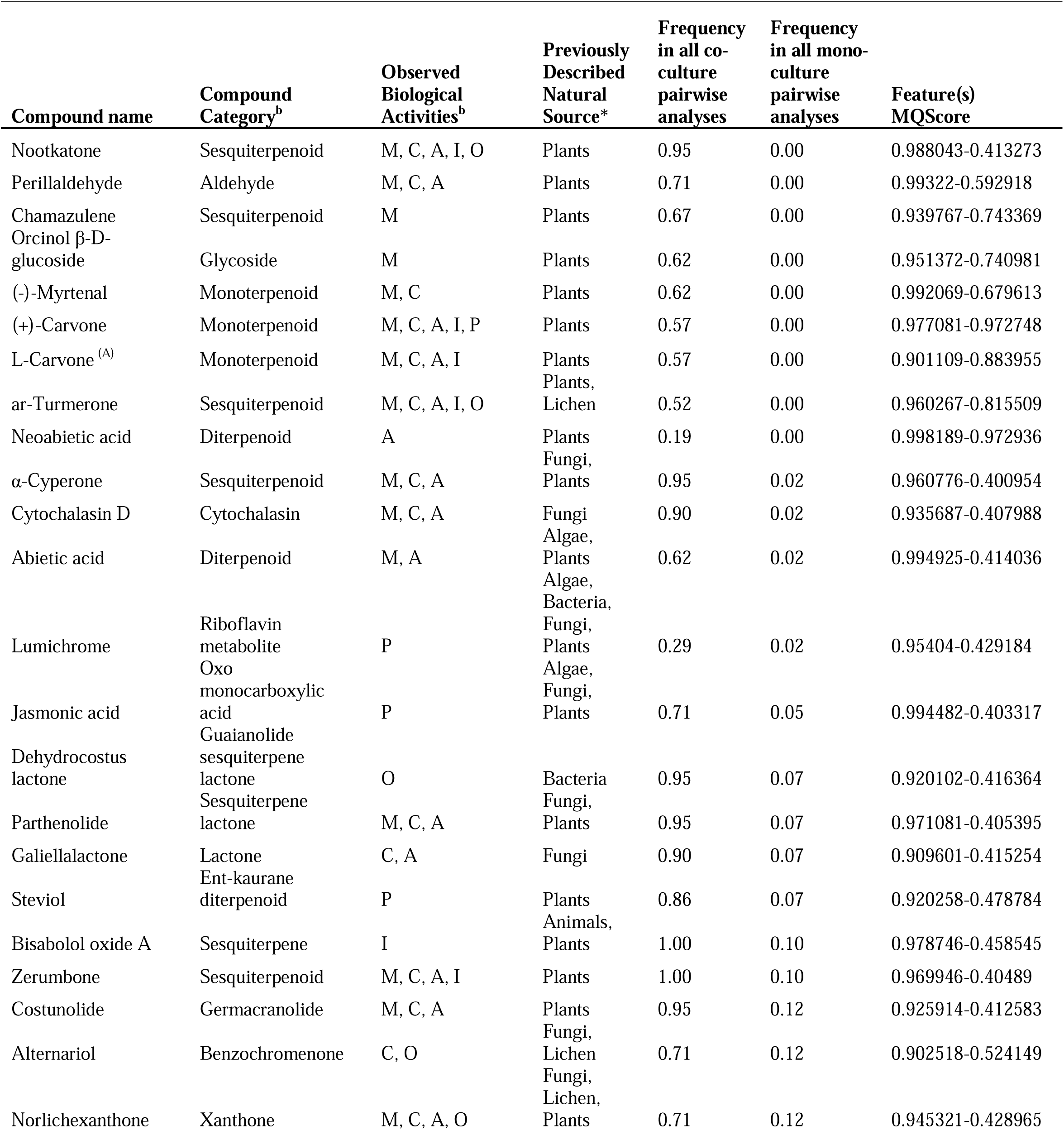
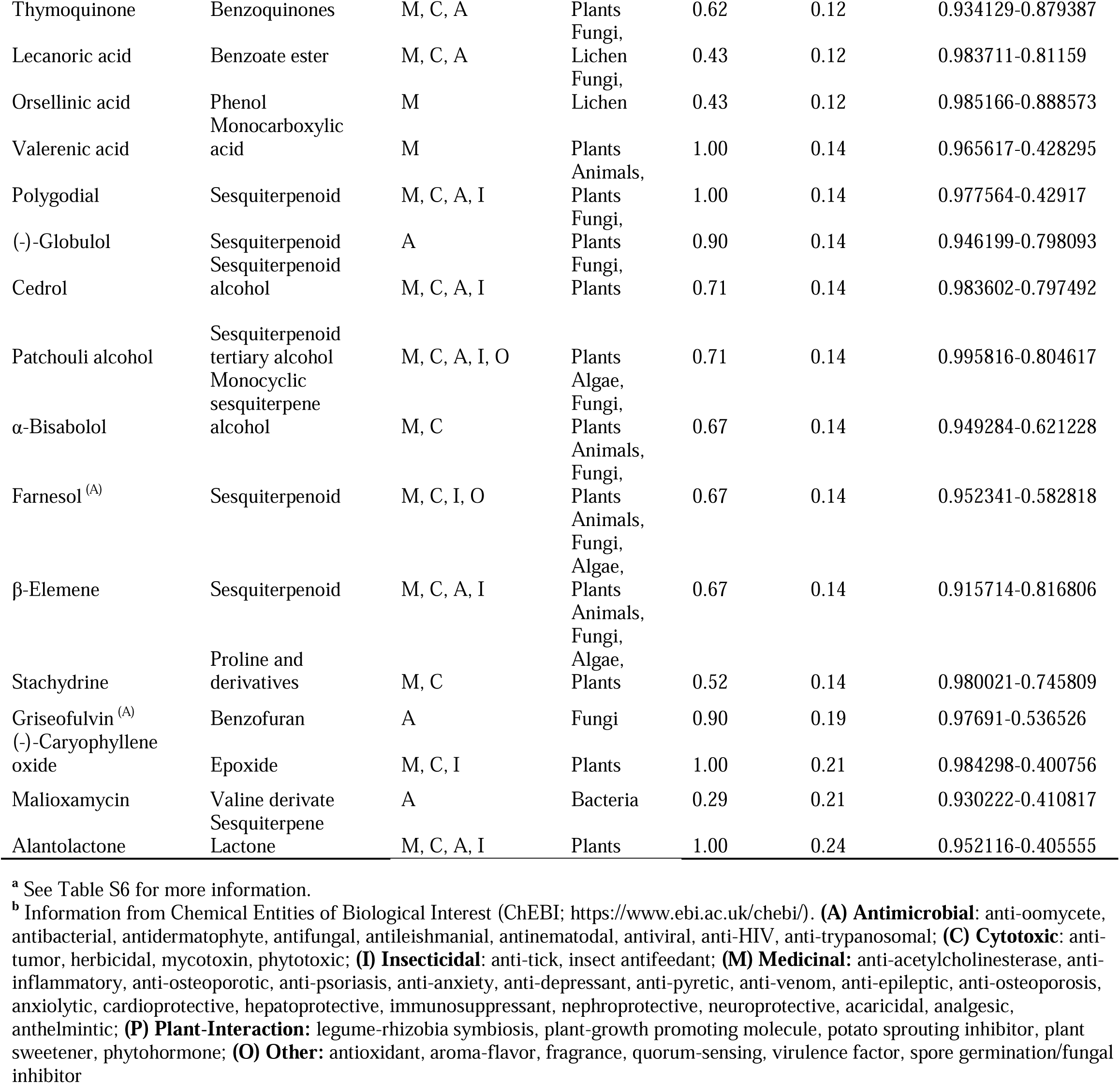
Subset of the putatively bioactive compounds produced by xylarialean fungi with their frequency in co-culture vs monoculture, and previouslyreported biological activities, uses, and natural sources^a^. Only compounds with MQScores >0.90 are shown.

Overall, we detected numerous features with putative matches to compounds typically identified in plants, including hormones, terpenes, and toxins. Features identified as the plant hormones jasmonic acid (JA) and abscisic acid (ABA) occurred in at least one co-culture of each of the seven strains (71% and 90% of all potential pairwise interactions) (Table S6). In general, features with matches to ABA were linked to a single unclustered gene in each of the genomes; however, in one case, the feature was linked to the ABA gene within a putative BGC in *P. punctata* (Table S5) (40). In at least one co-culture of each strain, we observed features with matches to the monoterpene carvone (Table 2), a component of dill and caraway essential oils with insecticidal and insect feeding-deterrent properties (40). Fungal features with hits to carvone were linked to numerous fungal genes, including some genes that reside within BGCs (Table S5).

Numerous features had hits to bioactive compounds with reported antimicrobial activity (Table 2, Table S6), such as the potent and non-toxic Gram-positive antibacterial compounds platencin and platensimycin from *Streptomyces platensis* (29, 41). Features identified as platencin were predominately found in co-cultures of *B. mediterranea*, whereas features linked to platensimycin were detected in all co-cultures of *X. flabelliformis* CBS 116.85 and *X. flabelliformis* NC1011. A feature identified as mitochondrial F1F0 ATP synthase inhibitor oligomycin B from *Streptomyces* spp. (42), occurred in four co-cultures involving *P. punctata*. A feature with hits to the antibiotic malioxamycin from *Streptomyces lydicu* (43) was exclusively detected in all co-cultures of *P. punctata*. Features matching the antibacterial and antifungal compound, asperlactone from *Aspergillus ochraceus* (44), were detected in all co-cultures of *B. mediterranea* and *N. diffusa*, as well as in co-cultures with *X. flabelliformis* CBS 123580. The broad-spectrum antibiotic 2,4-diacetylphloroglucinol (DAPG), which is considered the primary determinant of the biological control activity of *Pseudomonas fluorescens* F113 (45), was frequently observed in co-cultures with the three different strains of *X. flabelliformis*. Lastly, features matching the fungistatic compound griseofulvin and its structural analogs, dechlorogriseofulvin and griseofulvic acid (46) (Fig. S10), were observed in 90% of all co-culture combinations, as well as 19% of all monoculture comparisons, including monocultures of *B. mediterranea*, *P. punctata*, *X. arbuscula*, *X. flabelliformis* CBS 116.85, and *X. flabelliformis* NC1011 (Table 2; Fig. 5).

**Figure 5.**
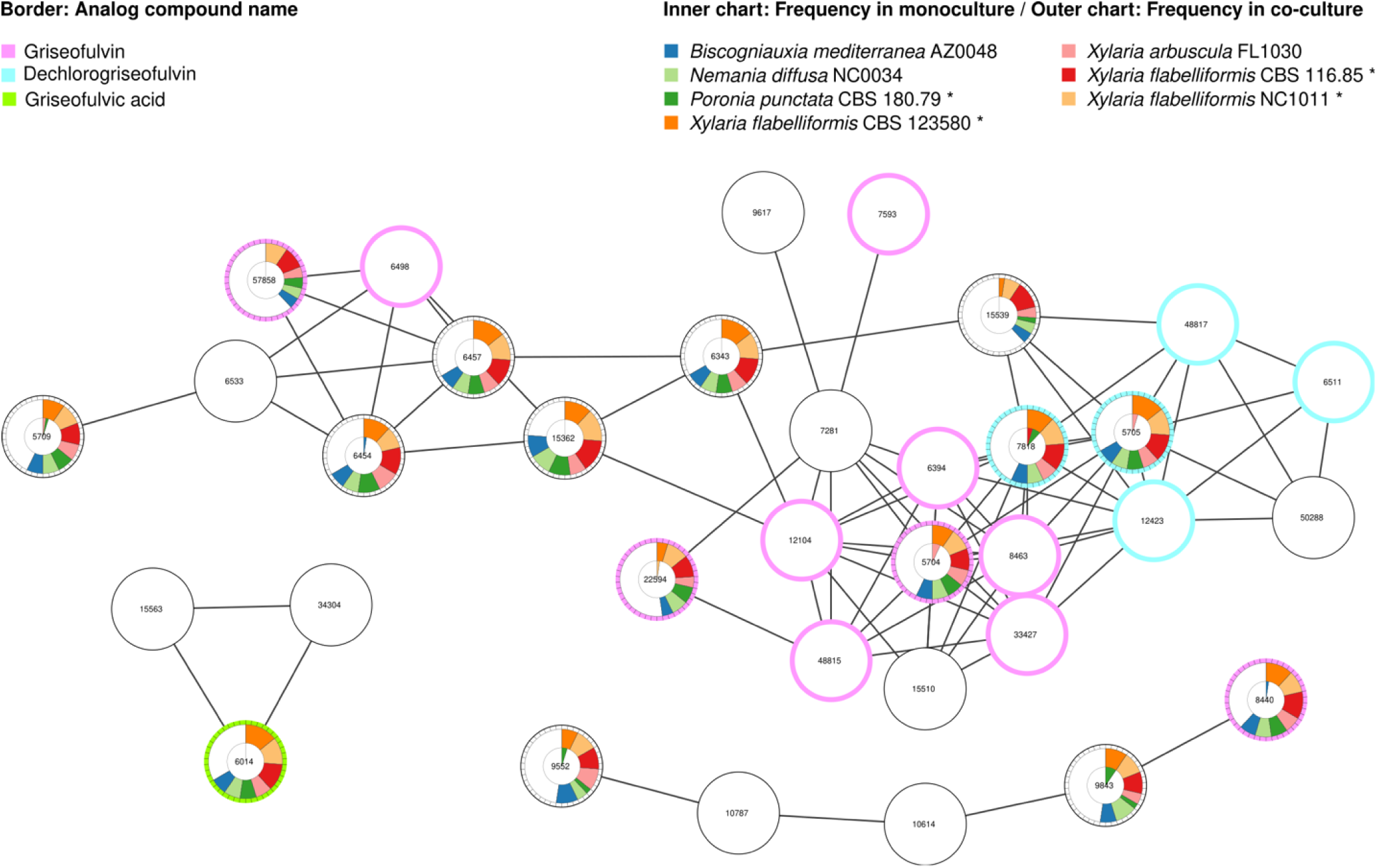
Compounds structurally related to griseofulvin were frequently detected among co-cultures yet rarely observed in monocultures. Network diagram showing features structurally related to griseofulvin. Each node represents a feature with its border colored by its analog compound identity: griseofulvin, dechlorogriseofulvin, or griseofulvic acid. Pie charts show the frequency with which each feature was detected as significantly more abundant in co-culture (outer chart) or in monoculture (inner chart) for each fungal strain (each shown in a different color). Asterisks (*) indicate strains with BGCs matching the griseofulvin biosynthetic gene cluster from *Penicillium aethiopicum* at the MIBiG repository (BGC0000070). Nodes were positioned by a pre-fuse force-directed layout algorithm, and edges between two nodes were weighted by the Cosine score. Ions were derived from the C18 chromatography and positive ionization mode. Numbers inside circles correspond to feature IDs.

## Discussion

Xylarialean fungi are of particular interest to biologists and chemists due to their bioactivity and chemical diversity (12, 13), which reflects a significantly higher richness and diversity of BGCs in xylarialean genomes compared to other filamentous fungi, including highly bioactive genera such as *Aspergillus* and *Penicillium* (11). Such enhanced secondary metabolism, which is linked to the number of HGT events and gene duplications (47), appeared greatest in the Xylariaceae *s.l.* clade that also has a greater ability to degrade lignocellulose as saprotrophs, as well as interact with a wide variety of plant and lichen hosts as endophytes compared to taxa in its sister clade, Hypoxylaceae (11).

Species that have evolved to occupy a specific ecological niche have also evolved to compete with one another, *sensu* Gause’s competitive exclusion principle (48). For example, the composition of fungal communities in decaying wood strongly reflects combative interactions between diverse basidiomycete fungi (49). In fungi, antagonistic interactions often occur primarily through the production of SM in the presence of the competing organism (23). Thus, to explain the diversification of BGCs in xylarialean fungi, we hypothesized that diverse competitive interactions—in both their free-living and symbiotic life stages—may exert selective pressure for both the horizontal transfer of BGCs and the maintenance of a diverse BGC repertoire (11). Here, we used co-culturing to test a series of hypotheses regarding how BGC content and phylogenetic identity might shape the diversity of metabolites produced by xylarialean fungi when grown alone or in competition with other strains.

Our first two hypotheses proposed that the richness of xylarialean metabolite production would be positively correlated with strain BGC content, and metabolite production in co-culture also would be impacted by the phylogenetic relationships among the strains. Specifically, we predicted that the three closely related strains of *X. flabelliformis—*whose genomes contain the highest number of predicted BGCs (11)*—*would be more likely to compete for resources due to shared niches compared to interactions between more distantly related organisms (48), and as a result, interactions between *X. flabelliformis* strains would produce the greatest number of metabolite features. However, for the seven xylarialean strains included here, there was no direct correlation between a strain’s BGC content and its metabolite production (in either monoculture or co-culture) (Pearson; P>0.05), as well as no clear relationship between the phylogenetic distance between competing strains and the richness of metabolite production. The lack of relationship between metabolite production and BGC content may be due to “silent” BGCs that were not activated under the conditions used here (i.e., ‘One Strain-Many Compounds’ (OSMAC) (50), or a non- 1:1 relationship between BGC number and SM number (e.g., multiple BGCs encode for a single SM (e.g., rhodochelin) or vice versa) (51). Additionally, although genomes of these strains are high-quality, the observed lack of correlation may reflect inaccurate BGC estimations due to incomplete/inaccurate assemblies or mis-annotations. Lastly, assaying a greater diversity of strains, including more distantly related strains, might reveal a pattern between metabolite production and BGC content and/or phylogenetic relationships.

Rather than closely related taxa producing the greatest numbers of metabolites, we found that interactions between distantly related *P. punctata* and either *X. flabelliformis* CBS116.94 or NC1011 resulted in the highest number of metabolites statistically over-abundant in co-culture occurred (n = 4,817 and 3,906 features, respectively). In general, all co-cultures with *P. punctata* as a partner typically resulted in a significantly higher mean number of over-abundant features, despite the fact that the genome of *P. punctata* has fewer predicted BGCs compared to the more closely related stains of *Xylaria* and *Nemania* (11). Surprisingly, despite no detectable growth on the media, we also observed that *P. punctata* could strongly inhibit the growth of *Nemania diffusa*, as well as three *X. flabelliformis* strains, presumably through the production of antimicrobial metabolites. In contrast to the lack of growth of *P. punctata*, one strain (*X. arbuscula*) overgrew its competitor in three of six interactions; however, overgrowth did not appear to substantially alter the number of features over-represented in *X. arbuscula* co-cultures compared to monocultures. Thus, our results suggest that xylarialean production of diverse metabolites in co-culture may not be easily predicted based the degree of fungal growth or inhibition/antagonism between strains in co-culture, phylogenetic distance between strains, or strain BGC content.

Our results instead suggest that the richness of metabolites produced by xylarialean fungi in co-culture may be strongly influenced by the ecological and habitat similarity of the competing strains. For example, six of the xylarialean strains included here were isolated as either endophytes of lichens/plants or saprotrophs of wood/bark and these species appear to be generalist saprotrophs based on the presence of identical or highly similar ITSnrDNA genotypes in decomposing plant materials (17), as well as a demonstrated capacity to degrade litter of *Quercus* and *Pinus* (11). In contrast, *P. punctata* has a distinct ecological niche as a specialist on herbivorous animal dung, which is a nutrient-rich, spatially limited, and ephemeral resource colonized by fungi, mycoparasites, bacteria, and invertebrates (52, 53). Reflecting its specialization on dung as a carbon source (54, 55) previous *in vitro* experiments found that *P. punctata* caused significantly lower mass loss on leaf litter of *Quercus* and *Pinus* compared to other strains of Xylariaceae *s.l.* (11). Thus, we posit that such ecological distinctiveness among competitors may have triggered the strong metabolic response observed in co-cultures with *P. punctata*. However, future studies that include a greater number and ecological diversity of strains are needed to better assess how fungal ecology, phylogenetic relationships, and genomic content influence metabolite production during competitive interactions.

Although fungal co-culturing is a long-established method to ‘turn on’ the synthesis of bioactive compounds that are typically not produced in pure culture (or only produced at low levels) (21, 23) and has led to the discovery of many novel fungal chemicals (23, 29, 56, 57), few studies to date have directly quantified the direct impact of co-culturing on metabolite production for a diverse set of strains. Here, our goal was to quantify the impact of co-culture on total metabolite richness, as well as determine how competitor identity influences xylarialean metabolome composition. Specifically, our last hypothesis was that co-culture interactions would result in a greater richness and diversity of metabolites being produced compared to monocultures, as the result of strain-specific metabolic responses to different competitors rather than the production of a uniform suite of metabolites for all competitors.

Consistent with previous studies and our hypothesis, xylarialean fungi produced significantly more metabolites in co-culture vs. monoculture (i.e., average of 2.2-fold increase). Furthermore, in support of our hypothesis that xylarialean fungi would exhibit a strain-specific response to competitors, we found that for each strain, each additional co-culture interaction resulted in an 11 to 14-fold increase in metabolite richness compared to the richness of metabolites with each additional monoculture. Thus, the high degree of metabolite diversity found here (i.e., over 9,000 features for only seven strains) is the result of largely unique metabolite responses in the majority of co-cultures: zero metabolite features were produced in all 21 co-culture combinations and <40% of features occurred in more than 10 different co-culture combinations (out of 21 possible).

However, one exception to this pattern was observed with *P. punctata*. Despite the higher number of over-abundant metabolites in pairs with *P. punctata,* interactions with the dung fungus produced the fewest number of unique features in co-culture: all *P. punctata* co-cultures contained the same 1,096 features regardless of the partner. In contrast, co-cultures of any strain with any of the three *X. flabelliformis* strains typically resulted in a large number of features unique to that interaction pair. The more uniform metabolite response of *P. punctata* to competitors vs. the unique responses of other xylarialean strains is congruent with our other results that suggest that ecological differences among xylarialean strains may influence their metabolite production in co-cultures more strongly than strain relatedness. These results are also consistent with the finding that xylarialean fungi that are ecological generalists have higher BGC diversity compared to strains that interact with fewer hosts and have a more limited ability to degrade lignocellulose (11).

Microorganisms typically employ secondary metabolites for niche exploitation, colonization, and competition with other microbes (5–7, 23, 58). Here, co-cultures of all strains resulted in a substantially higher number of features with hits to bioactive compounds with antiviral, bactericidal, fungicidal, insecticidal, herbicidal, and nematicidal activity (Table 2; Table S6). The observation that features related to broad-spectrum antimicrobial compounds were over-abundant in co-culture rather than monoculture is consistent with xylarialean fungi producing bioactive compounds as a mechanism to compete for resources within complex microbial communities often found in decomposing organic matter and soils (21, 58, 59). However, the production of antimicrobial or cytotoxic compounds by xylarialean fungi may also be related to their symbiotic interactions as endophytes.

For example, a previously noted positive relationship between the number of BGCs encoding non-ribosomal peptide synthases (NRPS) and endophytic host breadth for Xylariaceae *s.l.* taxa (11), provides evidence that xylarialean secondary metabolites have direct interactions with their host plants, similar to toxin production by fungal pathogens (60). However, endophytes can also produce antagonistic secondary metabolites that protect their hosts from pathogens and pests (7, 61). For example, the entomopathogenic endophyte *Beauveria bassiana* protects its hosts from insects (62). Here, in co-culture xylarialean fungi produced a metabolite feature similar to the insecticidal sesquiterpenoid nootkatone described from plants such as *Callitropsis nootkatensis* and grapefruit (63).

We found that xylarialean fungi also produced metabolite features putatively identified as steviol glycosides and the phytohormones ABA and JA, similar to previous research showing that numerous symbiotic fungi and bacteria can produce plant phytohormones that influence plant responses to abiotic stressors and modulate plant growth and development (64). For example, some strains of *Botrytis cinerea* can produce ABA (65). Likewise, certain fungi, such as *Lasiodiplodia theobromae*, can produce JA at high levels (66). Interestingly, here we found that xylarialean fungi produced ABA only in co-culture, which suggests a potential role for microbial interactions in stimulating the production of metabolites that could impact host plant fitness during symbiosis. Future studies are necessary to accurately identify these metabolites and determine their role in both competition and endophyte-host interactions.

### Importance of integrating genomic and metabolomic data

Although our untargeted approach was aimed at understanding phylogenetic and ecological factors that influence metabolite diversity in xylarialean fungi and does not provide the unambiguous identification of methods such as NMR (67), numerous features over-abundant in co-culture were also linked to genes or BGCs related to the production of secondary metabolites, providing support for putative compound matches. Our results also suggest that even fungi with significant genomic variation in BGC gene content and identity may be capable of producing structurally related compounds with similar antifungal activity. For example, features with matches to the fungistatic compound griseofulvin and its structural analogs (dechlorogriseofulvin and griseofulvic acid) were found in 90% of co-cultures and in monocultures of five xylarialean strains. However, genomes of only three of these five strains contain BGCs with MIBiG hits to griseofulvin (*X. flabelliformis* CBS 116.85, *X. flabelliformis* NC1011, and *P. punctata*) (11).

Thus, despite not identifying related BGCs in their genomes (11), we detected features related to griseofulvin in monocultures of *B. mediterranea* and *X. arbuscula* (11). However, it is possible these fungi may harbor BGCs for griseofulvin production but were undetected by bioinformatic methods due to fragmented assemblies or low similarity to the reference BGC from *Penicillium aethiopicum* IBT 5753 (MIBiG BGC0000070). For example, in genomes of *X. flabelliformis* and *P. punctata*, BGCs with matches to griseofulvin have very low similarity to the reference (i.e., 38% and 9% known cluster blast score, respectively), reflecting gene rearrangements and the loss of numerous *gsf* synthase and other genes compared to *P. aethiopicum* (11).

Although we observed griseofulvin production in some monocultures, it was primarily observed in co-cultures, consistent with griseofulvin’s fungistatic activity, whereby it inhibits but does not kill fungal competitors (68, 69). A previous co-culture study of an endophytic *X. flabelliformis* strain with *Aspergillus fischeri* also resulted in deadlock due to the production of griseofulvin (26). In that study, griseofulvin was completely extruded into the media in co-culture; yet in monoculture, the compound accumulated in outermost mycelia (26). Here, we detected griseofulvin in the media for monocultures of five of the seven strains, suggesting some fungal strains will secrete the compound even when grown alone.

Further targeted metabolomics research will be necessary to confirm that strains without a known griseofulvin BGC can produce the antifungal compound, as well as transcriptomics to identify the genes/BGCs that are upregulated during griseofulvin production. Overall, there remain significant challenges to link diverse xylarialean metabolites to BGCs as >90% of metabolite features we detected here were not identified at the SuperClass level and previously, only 25% of 6,879 putative BGCs predicted in 96 xylarialean genomes had hits to characterized BGCs in MIBiG (11).

### Conclusions

The diverse and strain-specific metabolites produced by generalist endophytic and saprotrophic xylarialean fungi in this study, along with the more uniform response of the specialist dung fungus to competitors, provide evidence that diverse competitive Interactions exert strong selection pressure for the diversification of biosynthetic gene clusters (BGCs) within the Xylariaceae *s.l.* clade. In addition to providing a better understanding of the evolutionary and ecological factors that influence the secondary metabolism of xylarialean fungi, our results also emphasize the incredible bioactivity of xylarialean fungi and their applications to the pharmaceutical, agricultural, food, and cosmetic industries as potential sources of putatively antiproliferative, anti-inflammatory, antioxidant, antiallergic, and sedative compounds, as well as mycotoxins, actin-binding cytochalasins, and phytohormones. Obtaining bioactive compounds is typically expensive and can have negative environmental impacts (70); thus, the potential for alternative fungal sources for some bioactive compounds would be both economically and environmentally beneficial.

Finally, the fact that most metabolite features detected here were unknown indicates the vast potential of xylarialean fungi towards the discovery of novel bioactive compounds. Future work will be necessary to accurately identify and characterize the structure of the diverse metabolites identified here, as well as experiments to validate metabolite bioactivity and identify potential ecological roles. Furthermore, to achieve a better understanding of the ecological and evolutionary drivers behind metabolite diversity in xylarialean fungi and other fungal clades, we must bridge the gap between fungal genetic potential and metabolic output via continued improvement of fungal metabolomic, genomic, and metagenomic data integration.

## Materials and Methods

### Fungal strain selection and co cultivation experiment

BGC content and phylogenetic position were used to select seven strains of Xylariaceae *s.l.* for co-culturing and metabolomics (11) (Fig. 1A; Supplemental Tables 7-9). A ∼6 mm diameter agar plug (removed from the growing edge of a 2-week-old pure culture of each strain using a sterile transfer tube) was placed on a 100 mm Petri dish with minimal media containing glucose either alone (i.e., monoculture) or in combination another strain (i.e., co-culture). Three replicate plates were inoculated for each monoculture and co-culture interaction. After one month at room temperature under ambient light/dark, defined interactions had occurred among the majority of fungal pairs and each plate was photographed (Supplemental Figs. 1-7). Each co-culture interaction was scored as either deadlock (i.e., visible inhibition zone) or invasion (i.e., one fungus grew over the other) following (49).

### Metabolite extraction and LC-MS/MS

As the majority of co-cultures had no mycelial contact, we harvested only media from the entire clear space between the hyphae of each strain (∼4-7 cm x 1 cm) using a sterile scalpel. In three cases, one strain overgrew the other, and we harvested a similar area of agar/mycelium in the interaction zone containing mycelia of both strains. For monocultures, a similar area of mycelium/agar was removed from the growing mycelial edge. A similar amount of media was also harvested from each negative control plate. Media/tissue was immediately flash-frozen in liquid N and stored at -80°C until lyophilized for 48 h. Freeze-dried samples were homogenized to a fine powder in a 2 ml tube containing stainless steel beads (Qiagen, Hilden, Germany) at 1,400 RPM for 30-sec intervals (maximum of 90 sec). Samples were extracted in MeOH and resuspended in methanol with an internal standard (ISTD) mix of isotopically labeled compounds (71), filtered, and analyzed on an Agilent 1290 UHPLC inline with a Thermo QExactive Orbitrap HF mass spectrometer (Thermo Scientific, San Jose, CA), implementing JGI LC-MS/MS ESI methods (71, 72). Normal phase and reverse chromatography were performed (3 uL injections) with randomized injection order. Blanks (methanol only) were run between each sample.

A list of features was generated with MZmine (73), removing isotopes, adducts, and features without MS/MS. The most intense fragmentation spectrum for each feature was uploaded to Global Natural Products Social Molecular Networking (GNPS) (74) and analyzed using the Feature-Based Molecular Network (FBMN) workflow. Peak heights were normalized using a global normalization approach adapted from NormalyzerDe (75). Only features that were significantly more abundant (log2FC = 2; adjusted P-value <0.05) in cultures vs. controls were kept for analyses (76). MetaboAnalyst 5.0 (77) was used to perform a data integrity check and data filtering using Interquartile Range (IQR). Features from each chromatography column and ionization mode were combined and compared between (i) each co-culture pair and their corresponding monocultures and (ii) all pairwise combinations of monocultures (see Fig. 1B). To connect features to putative genes/BGCs, MAGI 1.0 (30) was run with default setting using significant features and predicted genes of each strain (11).

## Data Availability

Raw metabolomics data is deposited in the Mass spectrometry Interactive Virtual Environment (http://massive.ucsd.edu/) with the accession number MSV000093277. For non-polar metabolites Feature Based Molecular Networking (FBMN) (results in positive and negative ionization modes (with minimum cosine score of 0.7 and at least 3 matching fragment ions for both library matches) are available at: https://gnps.ucsd.edu/ProteoSAFe/status.jsp?task=3b3f57d427b54d9fb5317bb94b73509b and https://gnps.ucsd.edu/ProteoSAFe/status.jsp?task=b3f3b9c0a43f46ceb140aa99ff2d0a20. For polar metabolites, FBMN results in positive and negative ionization modes are available at https://gnps.ucsd.edu/ProteoSAFe/status.jsp?task=e54e61e735f04bae9f55cd26cb25b7b5 and https://gnps.ucsd.edu/ProteoSAFe/status.jsp?task=ec1f5d74e6c349a284e69a1f3f897476. Genome data for all seven strains are available at JGI Mycocosm (https://mycocosm.jgi.doe.gov/mycocosm/home). Data files, scripts, and additional files are available at FigShare (https://doi.org/10.6084/m9.figshare.c.7539816).

## Acknowledgments

Funding for the project was provided by the DOE JGI Large-scale Community Science Project (Grant no. 503506 to JMU: https://doi.org/10.46936/10.25585/60001144) conducted by the U.S. Department of Energy Joint Genome Institute (https://ror.org/04xm1d337), a DOE Office of Science User Facility, that is supported by the Office of Science of the U.S. Department of Energy operated under Contract No. DE-AC02-05CH11231. MEEF was funded by the University of Arizona Office for Research, Innovation, and Impact and BIO5 Postdoctoral Fellowship Program. The authors thank Drs. M. Tfaily and J.H. Wisecaver for helpful discussion on the project; L.P. Moore for laboratory assistance; C. Orlando Ayala-Ortiz for sharing R scripts; and G. Sensibar and D. Odum at Tucson High Magnet School for generating the time-lapse of fungal interactions as part of their Southern Arizona Regional Science and Engineering Fair (SARSEF) project. The authors have no competing interests.

