## Supplementary Information for "Hyperdiverse, bioactive, and interaction-specific metabolites produced only in co-culture suggest diverse competitors may fuel secondary metabolism of xylarialean fungi"

### Supplemental Methods

***Fungal cultivation and media preparation.*** Fungi were grown on *Aspergillus* minimal media, with only essential compounds for fungal growth, as more rich media might impact the mass spectrometry results. *Aspergillus* minimal media containing a mixture of ~949 mL Nanopure water, 50 mL 20X Sodium nitrate salts, 1 mL 1000X Trace elements, 10 g D-glucose, and 10 g Agar (1). Media pH was adjusted to 6.5, sterilized by autoclaving for 30 minutes, and 15 mL was added into each Petri dish using a sterile serological pipette to ensure media uniformity among all replicate plates. The composition of 1 L of 20X Nitrate Salts consisted of 120 g Sodium Nitrate ($NaNO_3$), 10.4 g Potassium Chloride (KCl), 30.4 g Monopotassium phosphate ($KH_2PO_4$), 10.4 g Magnesium Sulfate Heptahydrate ($MgSO_4$-$7H_2O$) in Nanopure water. The composition of 100 mL of 1000X Trace Element solution consisted of 2.2 g Zinc Sulfate Heptahydrate ($ZnSO_4.7H_2O$), 1.1g Boric Acid ($H_3BO_3$), 0.5g Manganese Chloride ($MnCl_2$-$4H_2O$), 0.5g Ferrous Sulfate ($FeSO_4$-$7H_2O$), 0.17g Cobalt Chloride ($CoCl_2$-$6H_2O$), 0.16g Copper Sulfate ($CuSO_4$-$5H_2O$), 0.15g Sodium Molybdate ($NaMoO_4$-$2H_2O$), 5g Sodium EDTA ($Na_4EDTA$) in Nanopure water.

***Metabolite extraction***. To extract polar and non-polar metabolites, 500 µL of MeOH was added to each sample, briefly vortexed, sonicated for 5 minutes, and centrifuged for 5 min at 5000 rpm. The supernatant was removed, dried down in a SpeedVac (SPD111V, Thermo Scientific), and stored at -80 °C. Extraction controls were prepared in parallel using empty tubes. Dried extracts were then resuspended in 180 µL of methanol containing an internal standard (ISTD) mix of isotopically labeled compounds (see Table 11 in ref (2), but without 15N-guanine) and 2-Amino-3-Bromo-5-methylbenzoic acid (1 ug/mL, #R435902, Sigma), vortexed briefly, sonicated for 10 minutes and centrifuged for 5 minutes at 5000 rpm. After centrifugation, the resuspended extract was filtered through a 0.22 µm filter (UFC40GV0S, Millipore) for 2.5 min at 2500 rpm and transferred to a glass autosampler vial. Analyses were performed on an Agilent 1290 UHPLC inline with a Thermo QExactive Orbitrap HF (Thermo Scientific, San Jose, CA) mass spectrometer, implementing standard JGI LC-MS/MS ESI methods (2, 3). Both normal phase and reverse chromatography were performed on 3 uL injections, with MS1 collected in from mz range 80-1200 (reverse phase) or 70-1050 (normal phase) in both positive and negative polarity at 60k mass resolution, and MS2 fragmentation data acquired using stepped and averaged collision energies of 10, 20, 40 eV, or 20, 50, 60 eV (3rd replicate) at 17,500 resolution. Features with a retention time of < 0.7 minutes were included, although these accounted for only 1.5% of selected features. Across all 84 fungal samples and controls, liquid chromatography coupled with high-resolution tandem mass spectrometry (LC-MS/MS) detected 47,316 ions with unique mass-to-charge ratio (m/z) and retention time (RT) values (i.e., features) under both zwitterionic hydrophilic interaction LC (HILIC-Z) and C-18 reverse-phase columns with both positive and negative ionization mode.

***Data processing and statistical analysis.*** Among the 9,762 features significantly different from negative controls, a smaller number (959 HILIC-Z and 1,004 C-18) were assigned to 1,242 Simplified Molecular-Input Line-Entry System (SMILES) strings and 1,159 hashed International Chemical Identifier strings (i.e., InChIKey) (Table S3). In addition, 1,027 significant features had a mass spectrum that matched the GNPS database and could be assigned a feature identity based on the Feature-Based Molecular Networking (FBMN) workflow. Analyses were performed on data from each chromatography column and ionization mode separately, as well as with both columns and modes combined. After combining the data, we removed duplicate InCHEkeys, keeping only the feature with the best score. To preserve feature provenance, each feature in the combined set was renamed by appending the description of the corresponding chromatography column, ionization mode, row mass-to-charge ratio (m/z), and row retention time (RT) after the feature ID. We used custom R scripts and 'tidyverse' version 1.3.1 (4) in R (5) to remove features not significantly greater than controls (i.e., >log2fold). Proportional information on features by strain was summarized in a heatmap using the package ‘pheatmap’ v1.0.12(6) in R (5). The intersection of significantly more abundant features among different cultures was examined using the R packages 'ComplexUpset' version 1.3.3 (7), 'ComplexHeatmap' version 2.8.0 (8), and 'ggplot2' version 3.3.5 (9).

**List of Supplemental Tables (provided in separate Excel files)**

**Table S1.** Number of features detected and filtered for each chromatography column and ionization mode.

**Table S2.** Number of features that were significantly greater (i.e., log2FC, adjusted $P<0.05$) in either co-culture vs. monoculture for all 21 pairwise combinations. Features derived from both chromatographies and ionization modes were combined into a single dataset and analyzed together.

**Table S3.** Detailed information on features significantly more abundant in co-culture vs. monoculture comparisons, including hits to the Global Natural Products Social Molecular Networking (GNPS).

**Table S4.** Number of features and related compound gene-gene connections for both chromatography types (C18 and HILIC-Z) and ionization modes (positive and negative) combined and separate.

**Table S5.** Genes linked to compounds by MAGI using a 'MAGI score' ≥ 2, a 'reciprocal score' ≥ 1, and compound-gene connections labeled as 'direct', along with their functional annotation and prediction of HGT events derived from Franco et. al (2022).

**Table S6**. Subset of the putatively bioactive compounds, their frequency in xylarialean co-culture vs. monoculture, biological activities, uses, and previously reported biological sources.

**Table S7.** List of the seven Xylariaceae *sensu lato* strains used in this study, their ecological mode, substrate information, and the number of BGCs and KEGG pathways detected in their genomes (data from Franco et al. 2022).

**Table S8.** Number and type of Biosynthetic Gene Clusters (BGCs) predicted by antiSMASH v.5.1.0.

**Table S9.** Number of KEGG pathways identified in the genomes of the fungi used in this study. (A) The main pathway ("MainMap") is shown. (B) The main pathway ("MainMap") and the reference pathway ("SubMap") are shown.

**Supplemental Figures**

Focal isolate: *Biscogniauxia mediterranea* AZ0048

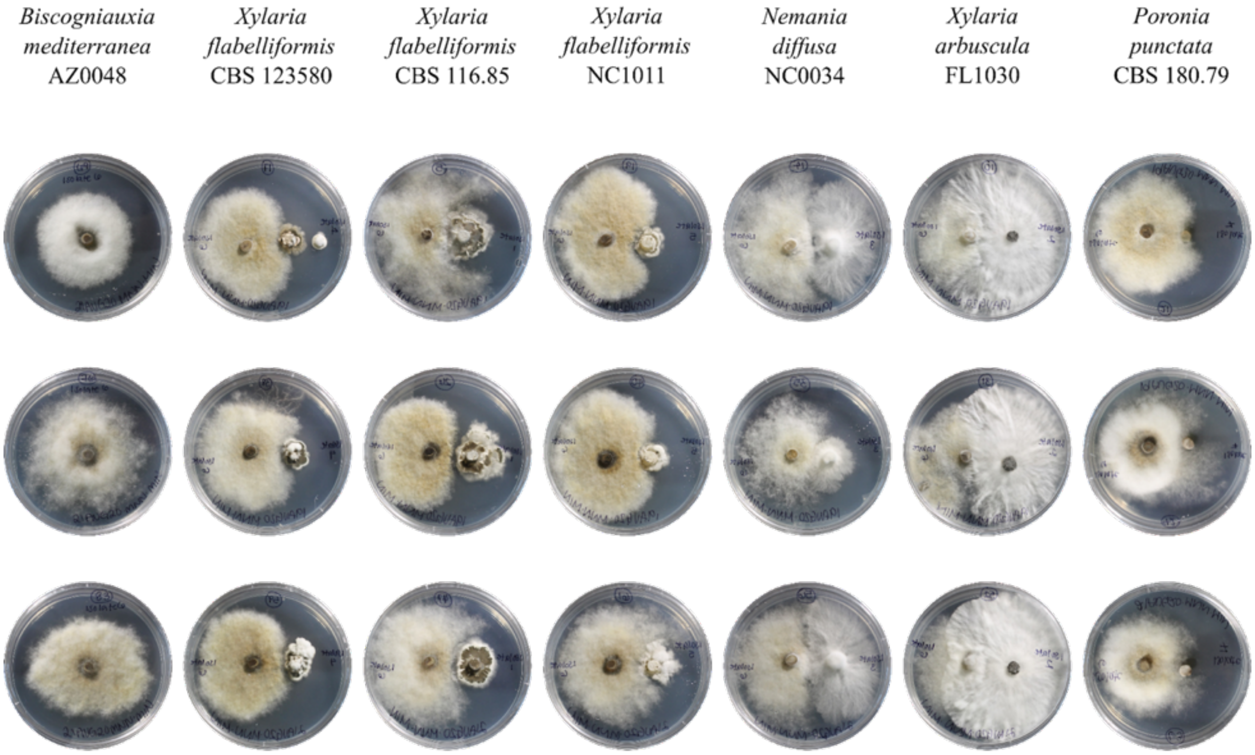


**Figure S1.** Monoculture and co-cultures of *Biscogniauxia mediterranea* AZ0048. Pictures correspond to cultures grown on *Aspergillus* minimal medium at room temperature, under ambient light/dark conditions, for one month.

Focal isolate: *Nemania diffusa* NC0034

| *Nemania diffusa* NC0034 | *Xylaria flabelliformis* CBS 123580 | *Xylaria flabelliformis* CBS 116.85 | *Xylaria flabelliformis* NC1011 | *Xylaria arbuscula* FL1030 | *Poronia punctata* CBS 180.79 | *Biscogniauxia mediterranea* AZ0048 |
|---|---|---|---|---|---|---|

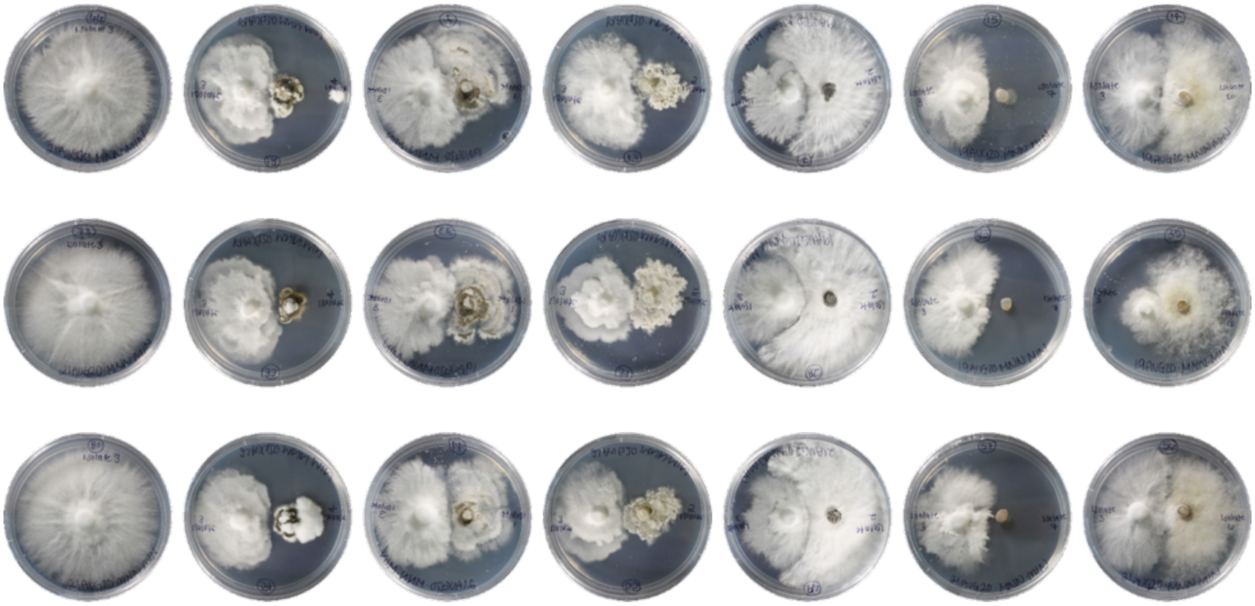

**Figure S2.** Monoculture and co-cultures of *Nemania diffusa* NC0034. Pictures correspond to cultures grown on *Aspergillus* minimal medium at room temperature, under ambient light/dark conditions, for one month.

Focal isolate: *Poronia punctata* CBS 180.79

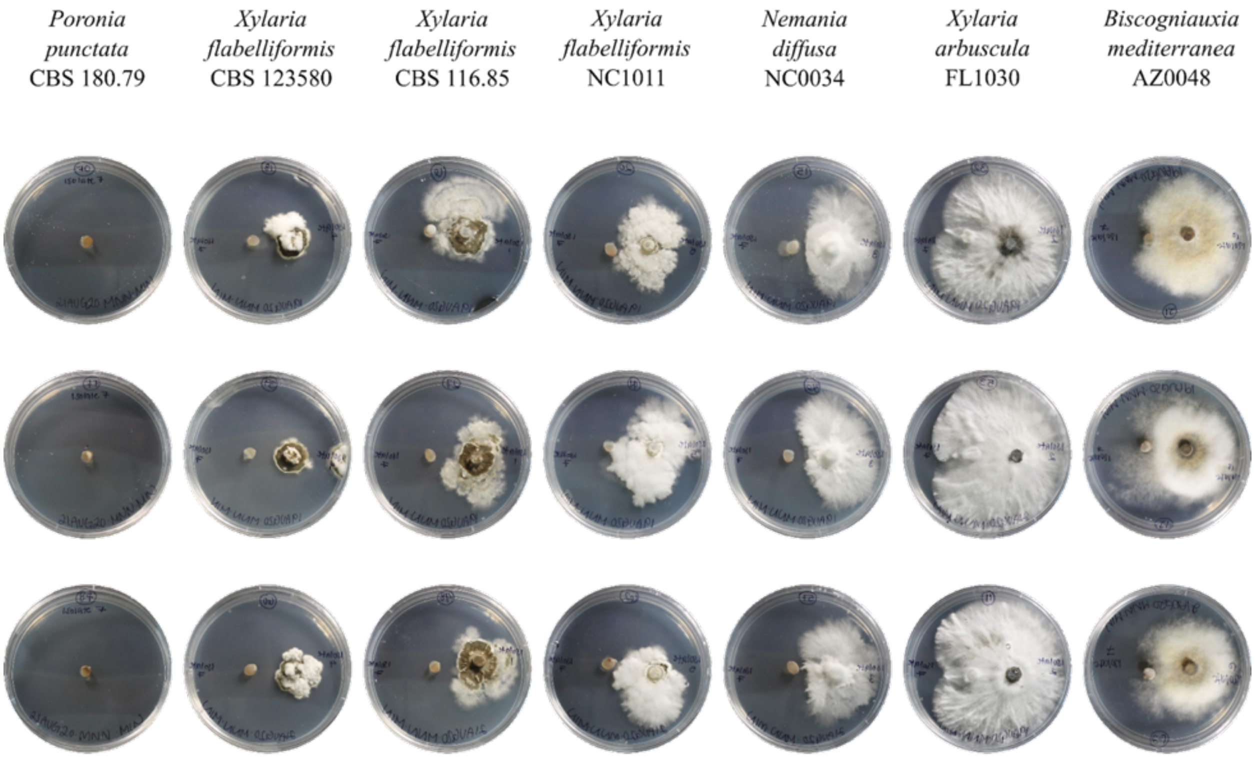


**Figure S3.** Monoculture and co-cultures of *Poronia punctata* CBS 180.79. Pictures correspond to cultures grown on *Aspergillus* minimal medium at room temperature, under ambient light/dark conditions, for one month.

Focal isolate: *Xylaria arbuscula* FL1030

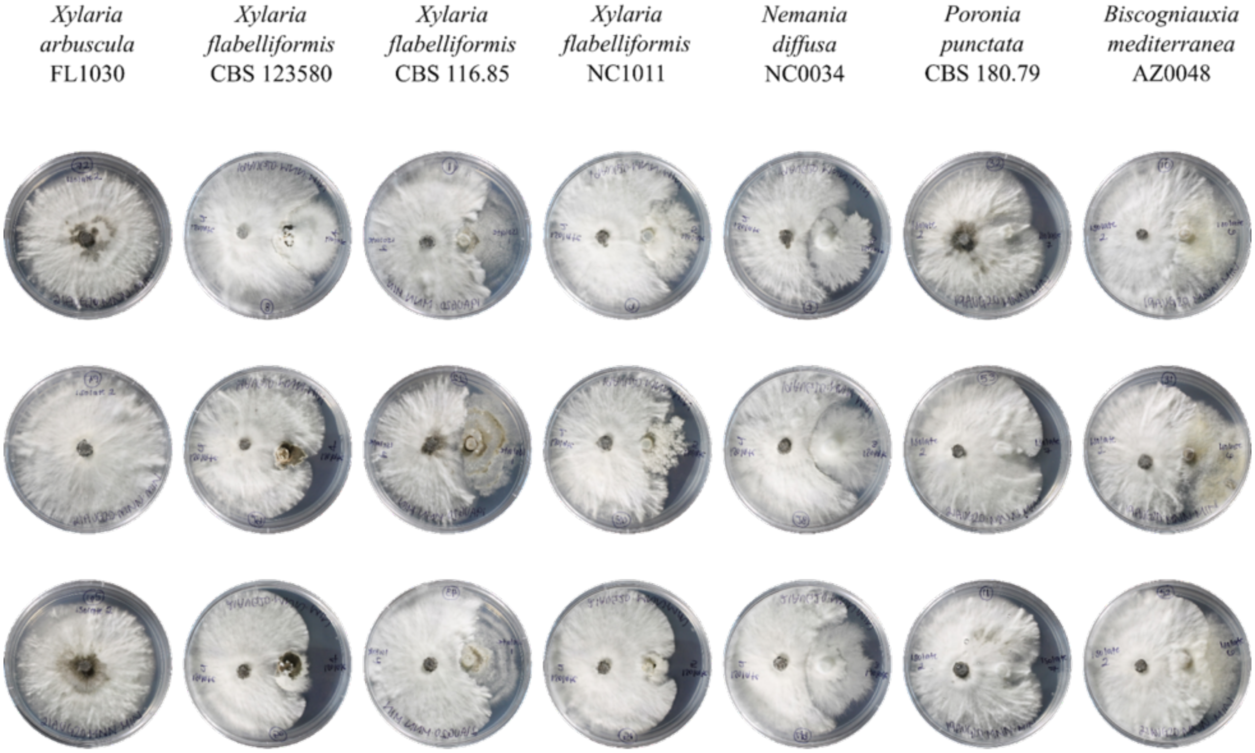


**Figure S4.** Monoculture and co-cultures of *Xylaria arbuscula* FL1030. Pictures correspond to cultures grown on *Aspergillus* minimal medium at room temperature, under ambient light/dark conditions, for one month.

Focal isolate: *Xylaria flabelliformis* CBS 116.85

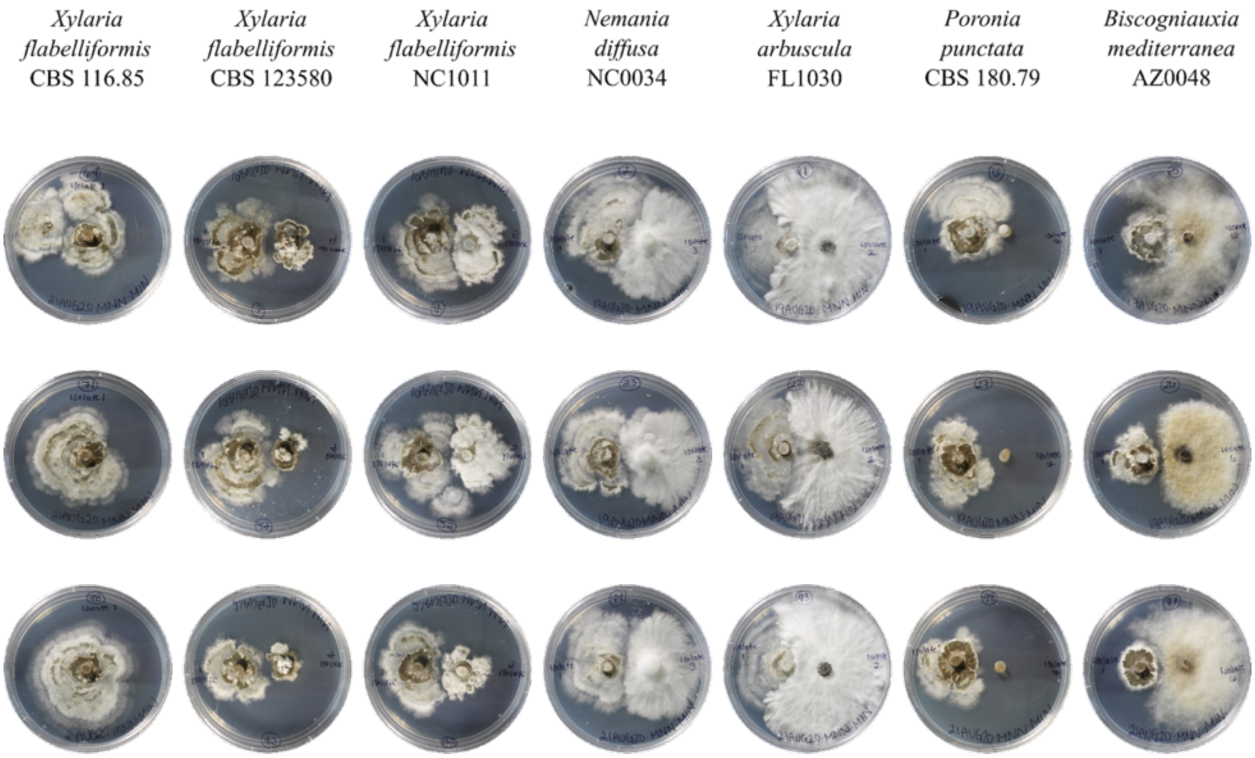


**Figure S5.** Monoculture and co-cultures of *Xylaria flabelliformis* CBS 116.85. Pictures correspond to cultures grown on *Aspergillus* minimal medium at room temperature, under ambient light/dark conditions, for one month.

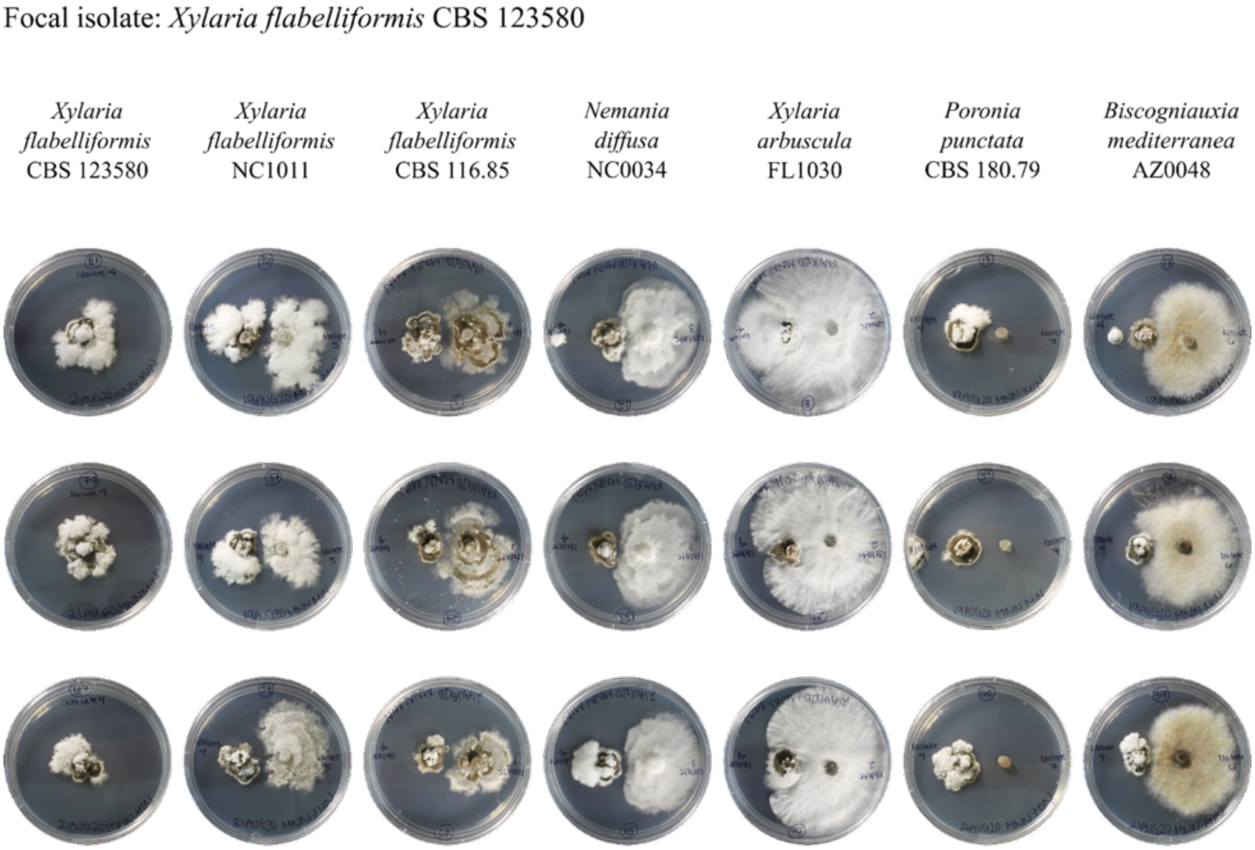


**Figure S6.** Monoculture and co-cultures of *Xylaria flabelliformis* CBS 123580. Pictures correspond to cultures grown on *Aspergillus* minimal medium at room temperature, under ambient light/dark conditions, for one month.

Focal isolate: *Xylaria flabelliformis* NC1011

| *Xylaria flabelliformis* NC1011 | *Xylaria flabelliformis* CBS 123580 | *Xylaria flabelliformis* CBS 116.85 | *Nemania diffusa* NC0034 | *Xylaria arbuscula* FL1030 | *Poronia punctata* CBS 180.79 | *Biscogniauxia mediterranea* AZ0048 |
|---|---|---|---|---|---|---|

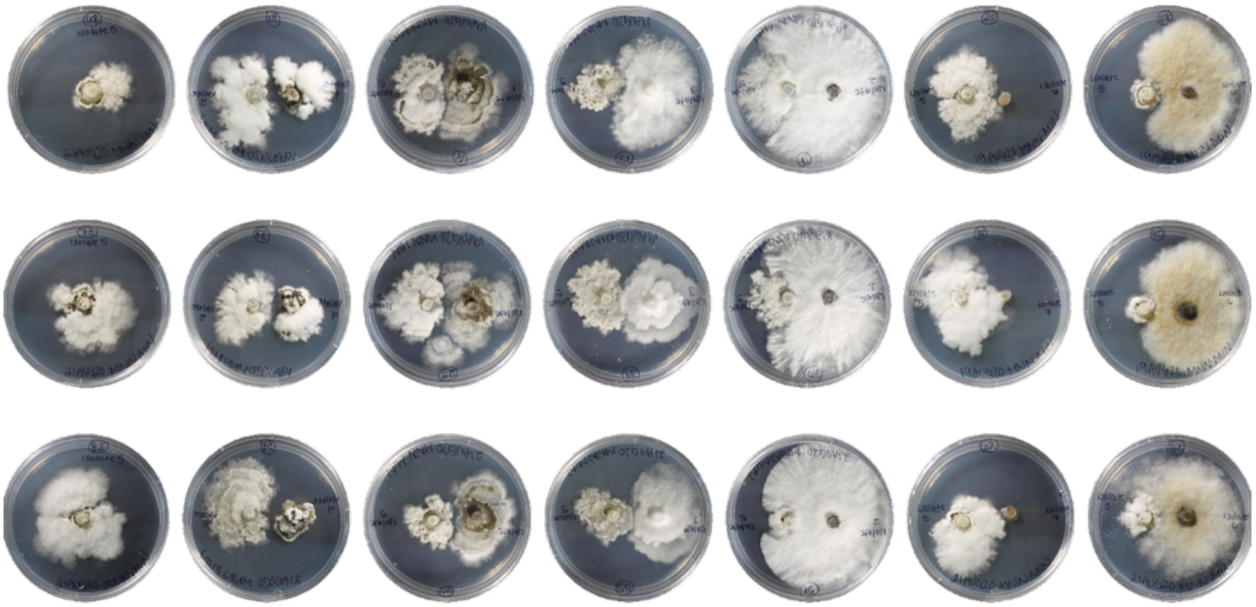

**Figure S7.** Monoculture and co-cultures of *Xylaria flabelliformis* NC1011. Pictures correspond to cultures grown on *Aspergillus* minimal medium at room temperature, under ambient light/dark conditions, for one month.

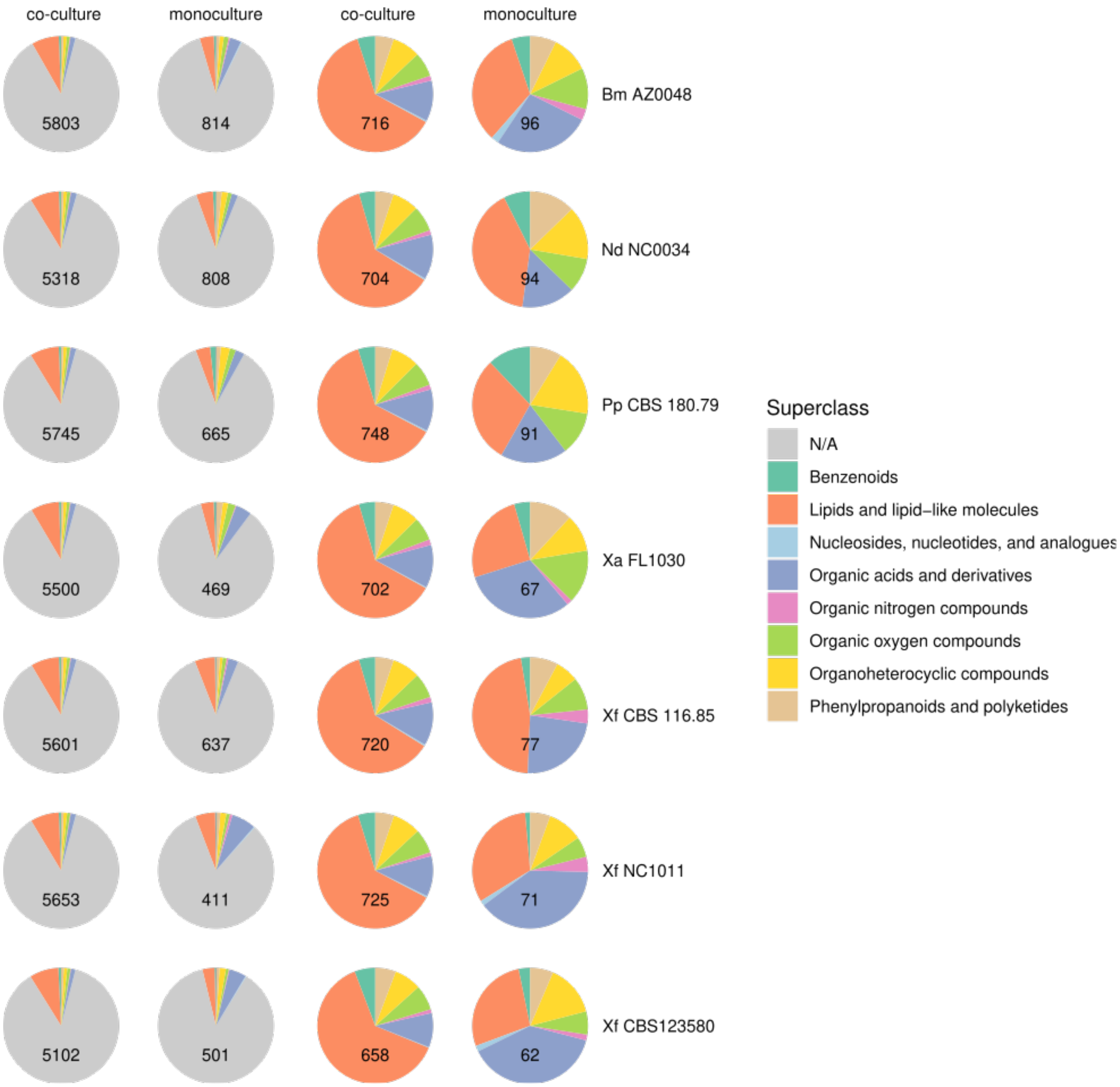


**Figure S8. Co-culturing altered metabolome composition compared to monocultures.** Pie charts represent the relative proportion of different metabolite superclasses for features over-represented exclusively in either co-culture or monocultures for each fungal strain. Pie charts in the two lefthand columns include features with no superclass identified (N/A), whereas pie charts in the two righthand columns only include features with a classification. The number of features included in each pie chart is shown. Classification is from the ClassyFire classification system. Features derived from both chromatography columns and ionization modes were combined into a single dataset and analyzed together. See Table S3 for detailed information on classification.

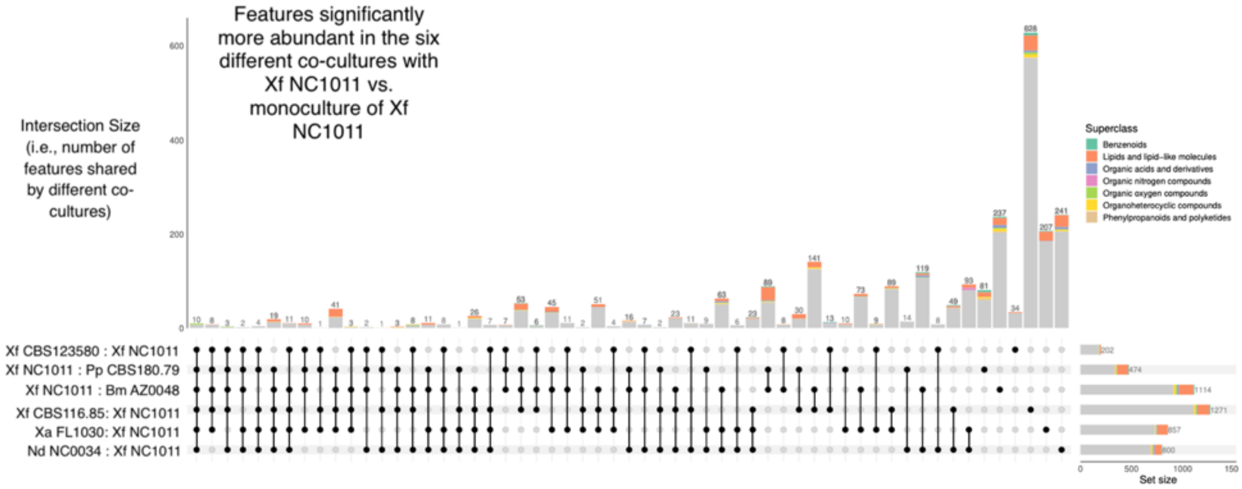
Features significantly more abundant in the six different co-cultures with Xf NC1011 vs. monoculture of Xf NC1011
Intersection Size (i.e., number of features shared by different co-cultures)
Superclass
Benzenoids
Lipids and lipid–like molecules
Organic acids and derivatives
Organic nitrogen compounds
Organic oxygen compounds
Organoheterocyclic compounds
Phenylpropanoids and polyketides
Xf CBS123580 : Xf NC1011
Xf NC1011 : Pp CBS180.79
Xf NC1011 : Bm AZ0048
Xf CBS116.85: Xf NC1011
Xa FL1030: Xf NC1011
Nd NC0034 : Xf NC1011
Set size

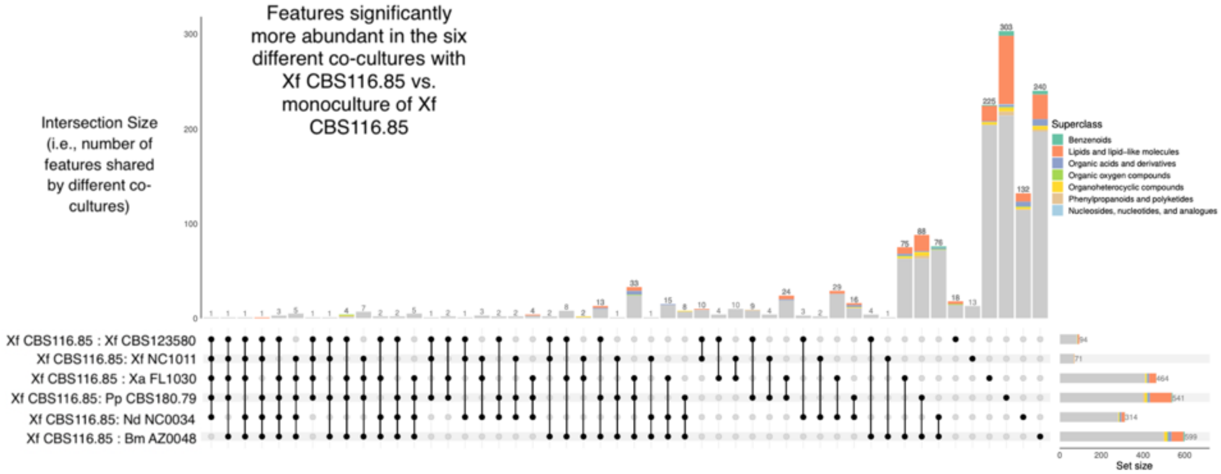
Features significantly more abundant in the six different co-cultures with Xf CBS116.85 vs. monoculture of Xf CBS116.85
Intersection Size (i.e., number of features shared by different co-cultures)
Superclass
Benzenoids
Lipids and lipid–like molecules
Organic acids and derivatives
Organic oxygen compounds
Organoheterocyclic compounds
Phenylpropanoids and polyketides
Nucleosides, nucleotides, and analogues
Xf CBS116.85 : Xf CBS123580
Xf CBS116.85: Xf NC1011
Xf CBS116.85 : Xa FL1030
Xf CBS116.85: Pp CBS180.79
Xf CBS116.85: Nd NC0034
Xf CBS116.85 : Bm AZ0048
Set size

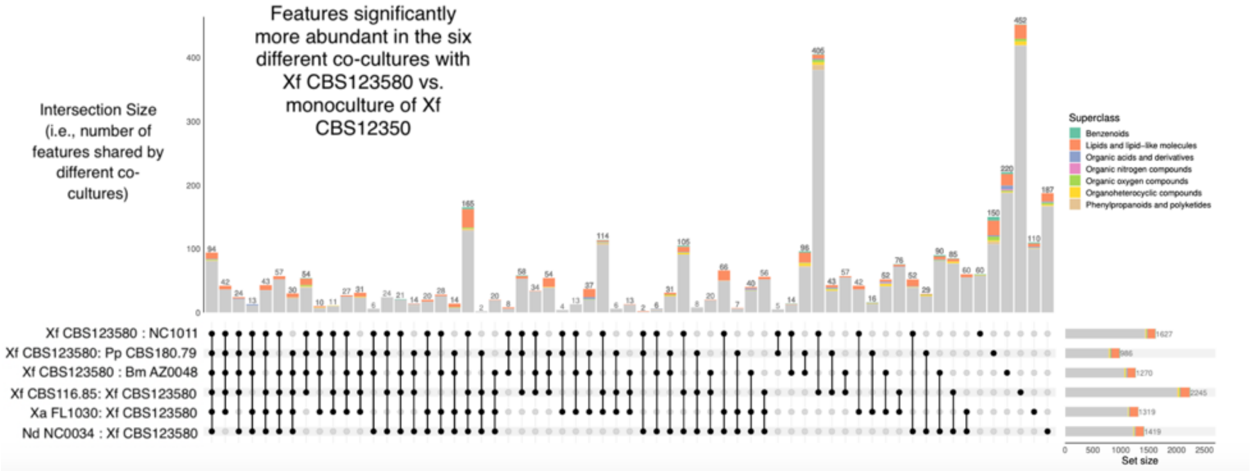
Features significantly more abundant in the six different co-cultures with Xf CBS123580 vs. monoculture of Xf CBS12350
Intersection Size (i.e., number of features shared by different co-cultures)
Superclass
Benzenoids
Lipids and lipid–like molecules
Organic acids and derivatives
Organic nitrogen compounds
Organic oxygen compounds
Organoheterocyclic compounds
Phenylpropanoids and polyketides
Xf CBS123580 : NC1011
Xf CBS123580: Pp CBS180.79
Xf CBS123580 : Bm AZ0048
Xf CBS116.85: Xf CBS123580
Xa FL1030: Xf CBS123580
Nd NC0034 : Xf CBS123580
Set size

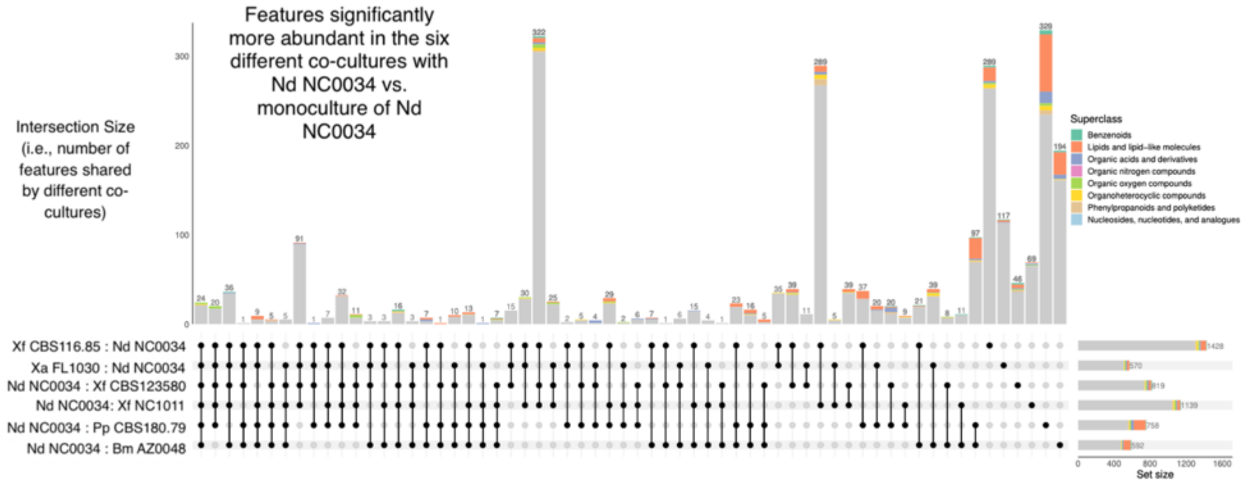
Features significantly more abundant in the six different co-cultures with Nd NC0034 vs. monoculture of Nd NC0034
Intersection Size (i.e., number of features shared by different co-cultures)
Superclass
Benzenoids
Lipids and lipid-like molecules
Organic acids and derivatives
Organic nitrogen compounds
Organic oxygen compounds
Organoheterocyclic compounds
Phenylpropanoids and polyketides
Nucleosides, nucleotides, and analogues
Xf CBS116.85 : Nd NC0034
Xa FL1030 : Nd NC0034
Nd NC0034 : Xf CBS123580
Nd NC0034: Xf NC1011
Nd NC0034 : Pp CBS180.79
Nd NC0034 : Bm AZ0048
Set size

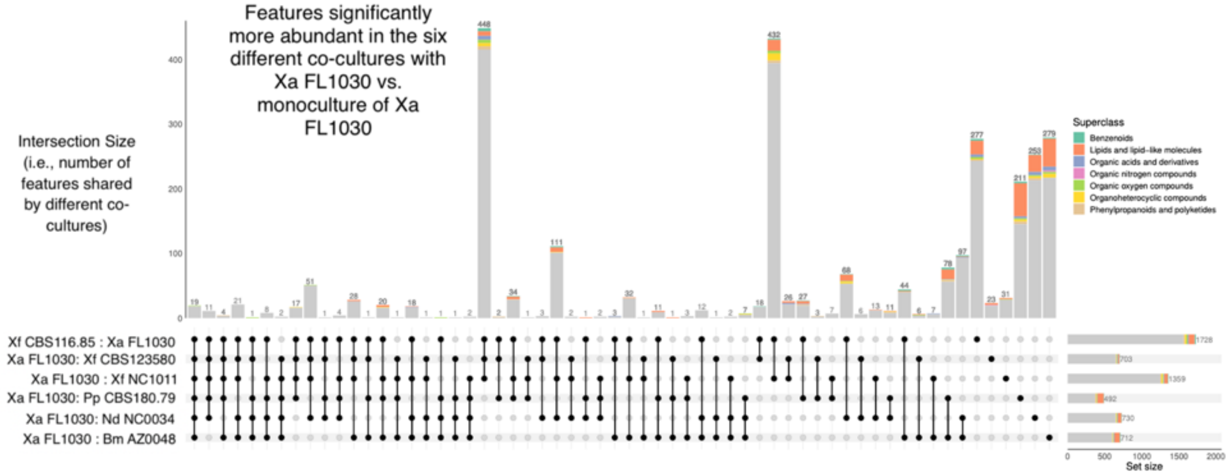
Features significantly more abundant in the six different co-cultures with Xa FL1030 vs. monoculture of Xa FL1030
Intersection Size (i.e., number of features shared by different co-cultures)
Superclass
Benzenoids
Lipids and lipid-like molecules
Organic acids and derivatives
Organic nitrogen compounds
Organic oxygen compounds
Organoheterocyclic compounds
Phenylpropanoids and polyketides
Xf CBS116.85 : Xa FL1030
Xa FL1030: Xf CBS123580
Xa FL1030 : Xf NC1011
Xa FL1030: Pp CBS180.79
Xa FL1030: Nd NC0034
Xa FL1030 : Bm AZ0048
Set size

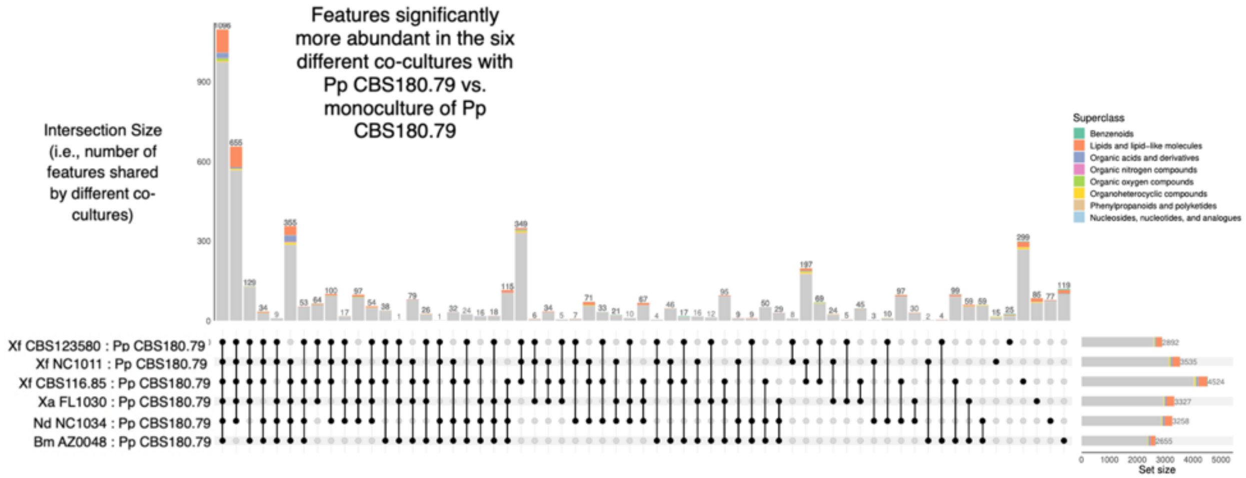
Features significantly more abundant in the six different co-cultures with Pp CBS180.79 vs. monoculture of Pp CBS180.79
Intersection Size (i.e., number of features shared by different co-cultures)
Superclass
Benzenoids
Lipids and lipid-like molecules
Organic acids and derivatives
Organic nitrogen compounds
Organic oxygen compounds
Organoheterocyclic compounds
Phenylpropanoids and polyketides
Nucleosides, nucleotides, and analogues
Xf CBS123580 : Pp CBS180.79
Xf NC1011 : Pp CBS180.79
Xf CBS116.85 : Pp CBS180.79
Xa FL1030 : Pp CBS180.79
Nd NC1034 : Pp CBS180.79
Bm AZ0048 : Pp CBS180.79
Set size

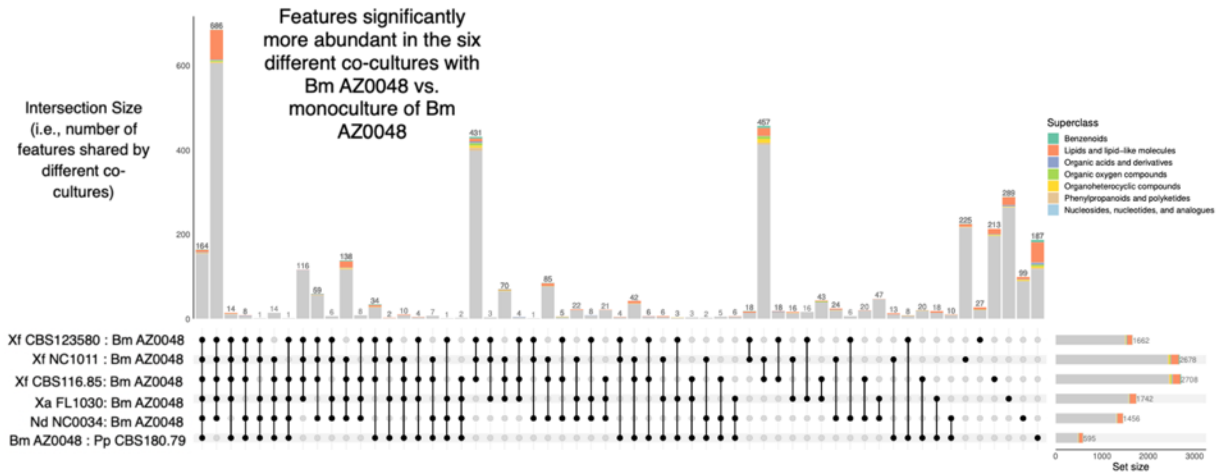


**Figure S9. Uniqueness of metabolite features among different co-culture combinations.** UpSet plots showing the overlap of features that were significantly more abundant in co-cultures compared to monoculture, as a function of different co-culture combinations. Each panel corresponds to one of the seven xylarialean strains. The height of each stacked bar indicates the number of features found (i.e., "intersection" size) across six different co-culture combinations compared to focal strain. For each intersection, dot color indicates whether a feature is present (black) or absent (grey) in each interaction pair. Black lines connect features in a set shared between >1 pair. Colors in each stacked bar correspond to feature classifications within each set at the 'Superclass level' using the ClassyFire (see legend). Horizontal stacked bars (i.e., "set" size) show the number of features significantly more abundant in the co-cultures of the focal strain compared to the monoculture. Features derived from both chromatography columns and ionization modes were combined into a single dataset for the analysis. Similar results are obtained when each column and ionization mode are analyzed separately. Fig. 5 shows feature overlap across 21 pairwise combinations.

**Griseofulvin**

Cosine similarity = 0.9531

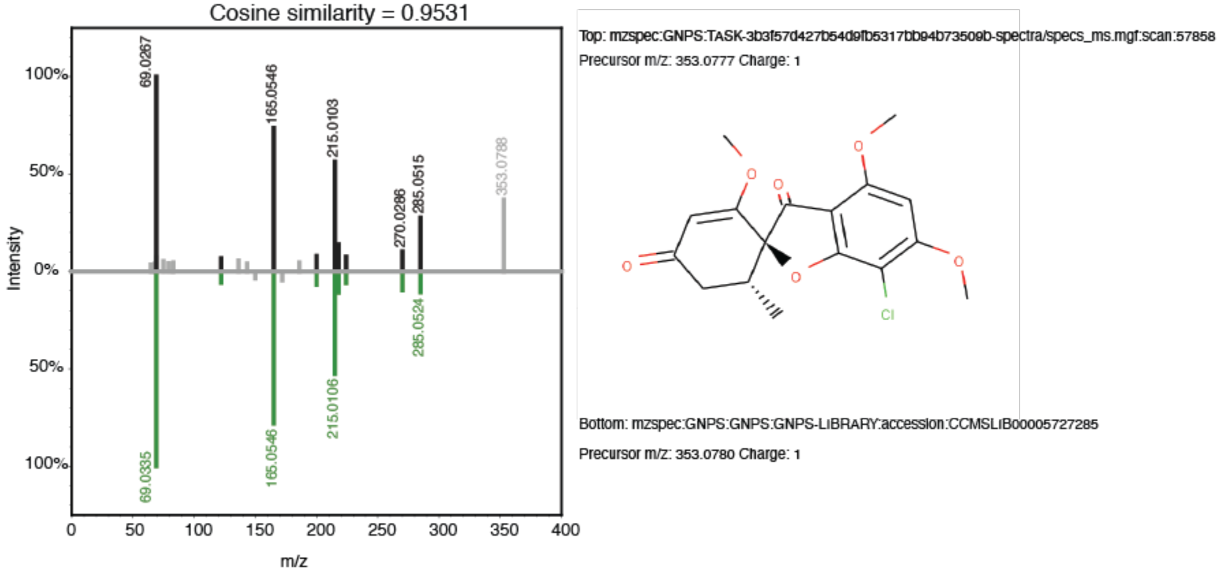


**Dechlorogriseofulvin**

Cosine similarity = 0.9113

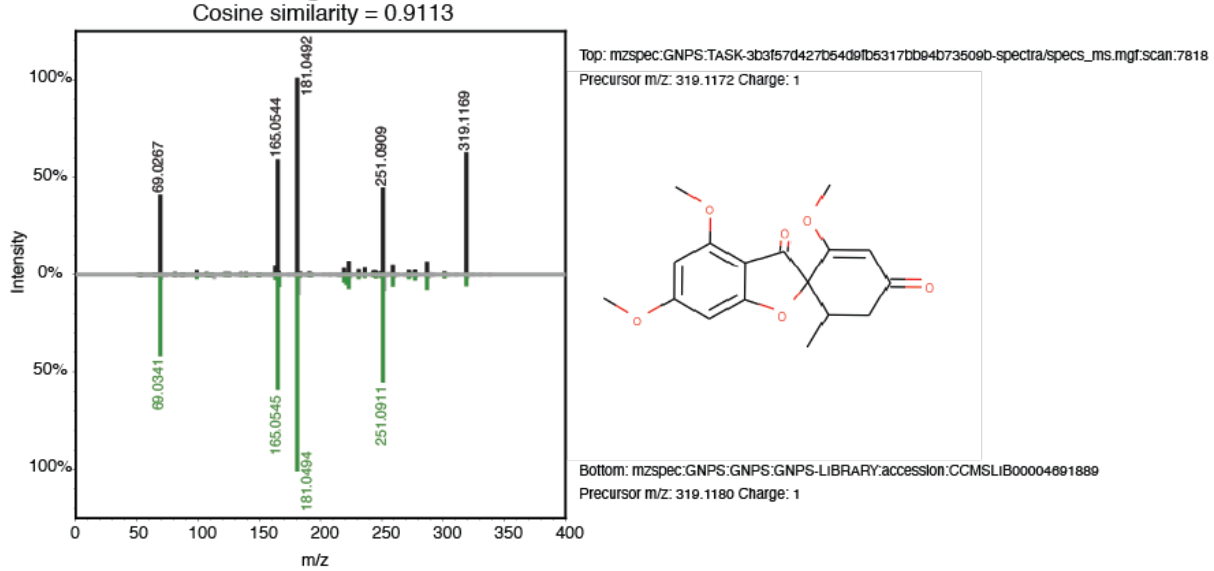


**Griseofulvic acid**

Cosine similarity = 0.3345

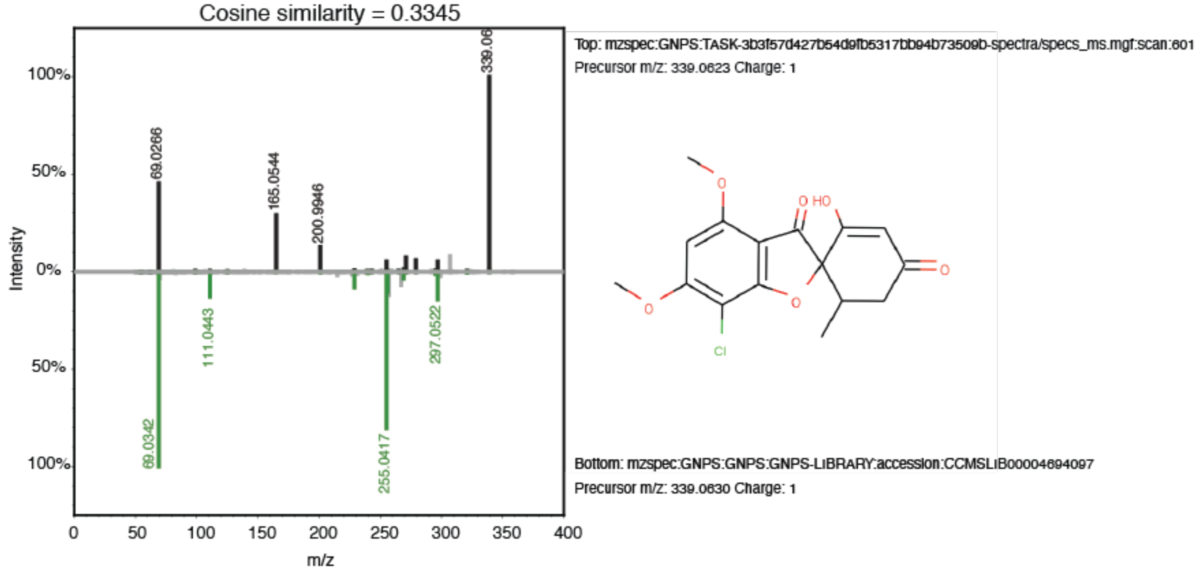

**Figure S10. Compounds structurally related to griseofulvin were frequently detected among xylarialean co-cultures** (see Fig. 6). Mirror plots between measured spectra and matching spectra from the Global Natural Product Social Molecular Networking (GNPS) library.

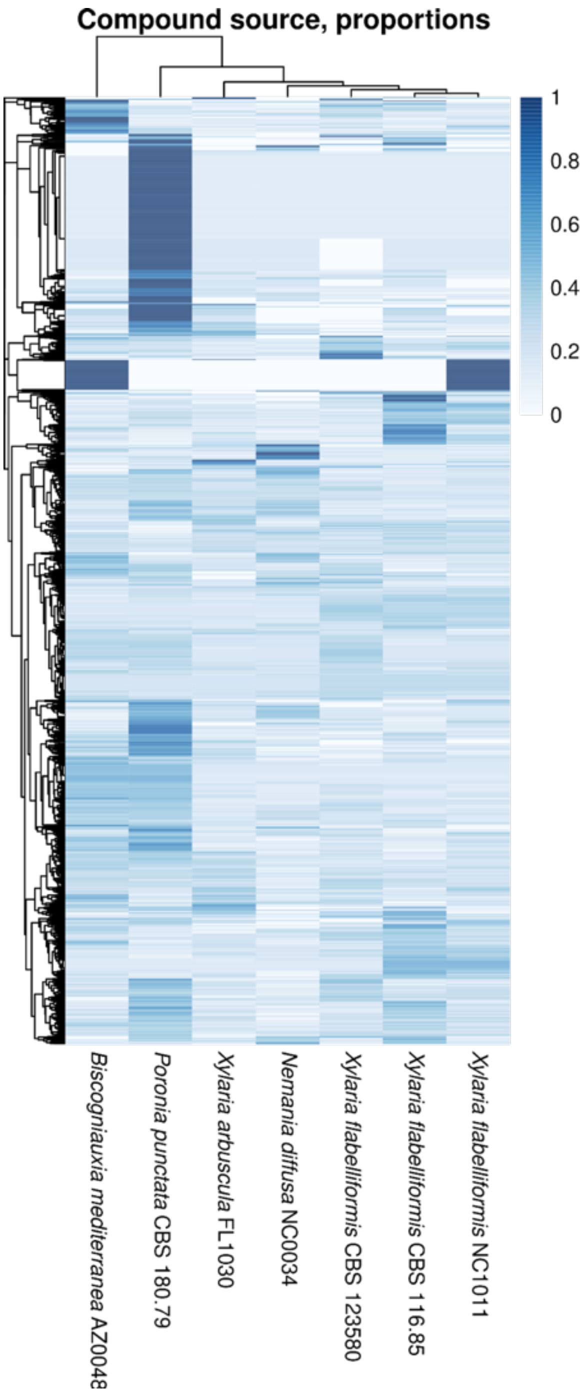


**Figure S11.** Heatmap based on the proportion of co-cultures (columns) where a feature (row) was found in co-culture for a particular strain (i.e., number of occurrences out of six different co-cultures). A proportion equal to 1.0 (dark blue) thus corresponds to compounds always detected in co-cultures with that strain, suggesting it may be the compound producer. Compounds and fungal strains were grouped by their phylogenetic relationships (shown in Fig. 1). Features derived from both chromatography columns and ionization modes were combined into a single dataset and analyzed together with duplicates removed. Features were grouped using hierarchical clustering with Euclidean distance and complete clustering in R (6).

**Additional Files:** https://doi.org/10.6084/m9.figshare.c.7539816
